# Reverse-Correlation Analysis of the Mechanosensation Circuit and Behavior in *C. elegans* Reveals Temporal and Spatial Encoding

**DOI:** 10.1101/147363

**Authors:** Daniel A. Porto, John Giblin, Yiran Zhao, Hang Lu

## Abstract

Animals must integrate the activity of multiple mechanoreceptors to navigate complex environments. In *Caenorhabditis elegans*, the general roles of the mechanosensory neurons have been defined, but most studies involve end-point or single-time-point measurements, and thus lack dynamical information. Here, we formulate a set of unbiased quantitative characterizations of the mechanosensory system by using reverse correlation analysis on behavior. We use a custom tracking, selective illumination, and optogenetics platform to compare two mechanosensory systems: the gentle-touch (TRNs) and harsh-touch (PVD) circuits. This method yields characteristic linear filters that allow for prediction of behavioral responses. The resulting filters are consistent with previous findings, and further provide new insights on the dynamics and spatial encoding of the systems. Our results suggest that the tiled network of the gentle-touch neurons has better resolution for spatial encoding than the harsh-touch neurons. Additionally, linear-nonlinear models can predict behavioral responses based only on sensory neuron activity. Our results capture the overall dynamics of behavior induced by the activation of sensory neurons, providing simple transformations that quantitatively characterize these systems. Furthermore, this platform can be extended to capture the behavioral dynamics induced by any neuron or other excitable cells in the animal.

## Introduction

A key function of the nervous system is to integrate the activity from a variety of sensory neurons and transform these neuronal signals into specific behavioral responses. This integration occurs not only across sensory modalities but also spatially and temporally within a single modality such as in mechanosensation ^1^. Characterizations of how the nervous system processes this information is vital for understanding brain function and allowing for prediction of behavioral responses. *Caenorhabditis elegans*, a nematode with a mapped connectome and powerful genetic and physiology tools, is an effective model organism for investigating relationships between sensory inputs and downstream activities ^2,3^. The components of the neural circuits involved in *C. elegans* mechanosensation have been elucidated through various genetic and behavioral analyses, coupled with neuronal cell ablation assays ^4–6^. Two sets of mechanoreceptors are specifically responsible for sensing touch throughout the body: the gentle touch sensing TRNs and harsh touch sensing PVDs ^7^. These specific neurons have been the focus of a number of studies, including genetic dissections of the mechanical signal transduction, their calcium responses and the eventual behavioral outcomes ^4,8–15^. However, most descriptions are specific to a single specific input stimulus, typically a single pulse with an eye lash or a metal pick, and a single behavioral output. This leaves unexplored space of the stimuli and outputs, leading to descriptions that are potentially biased toward a specific stimulus, and not allowing for the generalizable prediction of the system.

To map the transformations between mechanoreceptor neurons and behavioral outputs, we sought to model these transformations in an unbiased quantitative framework that captures the systems’ dynamics in a predictive manner. This is computationally challenging because of the stochasticity and complexity of the animal’s behavioral repertoire, as well as the various time scales and frequencies relevant in the system^16–18^. A successful technique for characterizing neuronal systems is the use of reverse correlation analysis with a white noise stimulus ^19–26^. This methodology is commonly applied in sensory physiology to model a sensory neuron’s response to natural stimuli as a linear filter. The computed linear filters provide a complete description of the linear dynamics of the neuron, and can be used in conjunction with a nonlinear filter to accurately model its function ^21,27–30^. This technique has also been extended to modeling sensory neurons^31^ and behavior in invertebrates^32–36^. However, this technique has not been extended to model and contrast the spatial and temporal properties of behavioral responses to the gentle and harsh touch mechanosensory neurons.

Although reverse correlation analysis allows for accurate estimations of system dynamics, several experimental obstacles hinder its applicability to the mechanosensory circuits in *C. elegans* at present. Current techniques for delivering precise mechanical stimuli to animals involve the delivery of a mechanical force via a stylus or microfluidic device to specific locations on the animal’s body ^9,14,15,37^. Although ideal for neuronal imaging, these techniques require the immobilization of animals with glue or other techniques, and therefore, do not allow for reverse correlation analysis with behavior response dynamics. Additionally, many of these techniques have a low experimental throughput, and cannot provide the large sample sizes required for reverse correlation studies. One technique that overcomes these challenges is to couple optogenetics with behavior, as a light stimulus is more easily controlled, and can be used to activate specific neurons in freely moving animals ^34,35,38^. This fictive stimulus has the added benefit of bypassing differences in native receptor protein expressions, allowing for comparison between sensory systems. In order to apply light stimuli with spatial resolution to activate specific regions of sensory neurons, we adapted a previously developed tracking platform with selective illumination ^39^. The custom microscopy system uses a projector and computer vision tools to track the animal, allowing for the delivery of spatially and temporally resolved stimuli required for white noise signal delivery.

Combining these tools, we developed an experimental and computational pipeline for performing white noise analysis on *C. elegans*, and apply this method to elucidate models of transformations between mechanosensory neuron activity and behavioral response. Using our platform, we computed linear filters that characterize the dynamics of the gentle touch sensing TRNs and harsh touch sensing PVDs. These filters provide a quantitative framework for the functions of these neurons, and allowed for the investigation of differences in spatial encoding. Furthermore, this method allowed us to create models that accurately predict behavioral changes in response to mechanosensory neuron activity. Our method provides simple transformations that quantitatively characterize these systems by capturing the spatiotemporal dynamics of behaviors induced by optogenetic activation of sensory neurons.

## Results

### Reverse-correlation analysis using optogenetics and behavior tracking

To illuminate the differences between the mechanosensory systems, we characterize and compare the dynamics for these two anatomically distinct sets of mechanosensory neurons: the gentle touch sensing TRNs and the harsh touch sensing PVDs (Fig 1A). In order to use reverse correlation for modeling behavioral responses, the two main experimental requirements are the delivery of a white noise stimulus and accurate measurements of the output. For the stimulus, we used optogenetics to directly activate the mechanosensory neurons with a white noise signal. This unmediated input enabled us to activate neurons regardless of expression of mechanotransductive channels. This allows the comparison of how the two systems and their morphologies control downstream activity, rather than differences in their sensory activation. Additionally, whereas a natural stimulus can activate additional sensory neurons and possibly interfere with the characterization, the optogenetic stimulus will only activate the neurons expressing channelrhodopsin. Therefore, the resulting filters characterize the dynamics of behaviors exclusively in response to activation of specific sensory neurons. Our tracking platform ^39^ enables the delivery of patterned illumination while simultaneously tracking individual animals, allowing for selective activation of specific sections of transgenic animals with high spatial and temporal precision (Fig 1B, Movie S1, Methods). We used this platform to deliver the white noise light stimulus for reverse correlation; we activate mechanosensory neurons with a pseudo-random m-sequence pattern, a spectrally unbiased binary signal (Methods).

**Fig 1:**
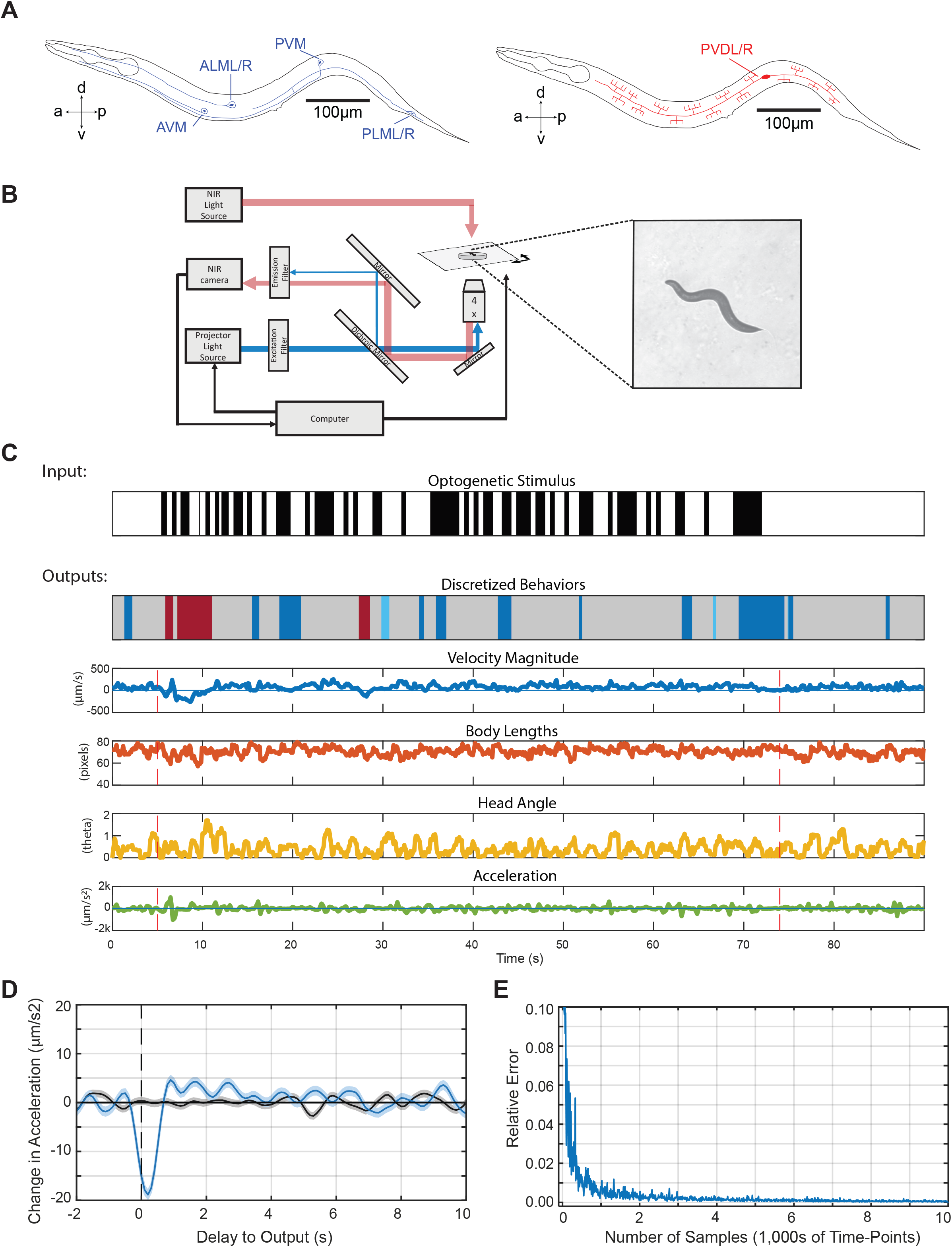
Reverse correlation analysis of mechanosensory neurons enabled by tracking and selective illumination platform. (A) Mechanosensory neurons characterized in this study. The gentle touch sensing neurons ALML/R, AVM, PVM, and PLML/R (blue) and harsh touch sensing neurons PVDL/R (red). (B) Schematic of custom tracking system with selective illumination used for reverse correlation experiments (Methods). A projector is used as the light source to enable selective illumination. Captured video frames are processed in real-time to deliver accurate light patterns on moving animals. (C) Sample stimulus and extracted quantified behavior traces obtained from the custom platform and analysis script (Methods). Input is a binary signal of On and Off. Outputs are characterized for both “discrete” and “continuous” behaviors. Discretized behaviors are classified based on a custom behavior analysis script (Methods). Colors represented in sample output: dark blue represents a pause, red represent reversals, light blue represents turns. (D) A sample filter computed using the BWA computation (Acceleration Response to Anterior TRN, n = 88,031 time-points). (E) The speed of convergence for the BWA as a function of the amount of data used to train the model. The error converges to a relative tolerance of δ<0.005 after 30,000 time-points.

The outputs we seek to characterize are the behavioral responses of animals using the optogenetic stimuli as inputs. We developed a custom computer vision algorithm (Methods) to analyze recordings of animals’ behavior in a high-throughput and unbiased manner. The worm’s posture and position are extracted for each frame, which are then used to quantify various “continuous” behaviors such as instantaneous velocity, instantaneous head angle, and instantaneous acceleration (Fig 1C). In addition to these “continuous” behaviors we also quantified and categorized several classical “discrete” behaviors such as reversals, pauses, and omega turns ^18,40–42^ (Fig 1C, Methods). Each of these continuous and discrete variables was used as a separate output for reverse correlation analysis, yielding a filter that can be used to predict behavior responses to any arbitrary stimulus patterns. By using filters for a large portion of the worm’s behavioral repertoire, we can describe the overall behavioral response when stimulating specific mechanosensory neurons.

Using the white noise light stimulus for optogenetics and the quantified behavioral responses, we next apply reverse correlation to model *C. elegans* response as transformations of linear and non-linear filters. Classically, when characterizing mammalian neuronal systems, a neuron’s response is modeled by computing the average of the stimuli that preceded its action potentials (spike-triggered average or STA) or its subthreshold voltages (voltage-weighted average or VWA) ^29^. Analogously, we estimate the dynamics of *C. elegans* response by computing the behavior weighted average of the stimulus (BWA). When stimulating specific segments of the mechanosensory systems, the BWA represents how the animals characteristically transform patterns of activity of those neurons into specific behaviors, providing a filter estimation of this transformation (Fig 1D).

In order to accurately estimate these linear filters, a large sample size is required to test enough input values ^20,30^. To estimate the number of samples required in our system, we characterized the speed of convergence of computed filters as the number of input samples increased (Movie S2). We characterized the convergence of filters by computing the L2 norm of the difference between subsequent filters (computed as the absolute difference between filters). We found that our system converges (to a relative tolerance of δ<0.005) after using roughly 30,000 frames of tracking data (Fig 1E). With our experimental conditions, this is equivalent to a sample size of roughly 30 animals (Methods).

### Linear Filters for anterior and posterior touch receptor neurons (TRNs) robustly capture behavioral dynamics

We first used our method to characterize responses to the touch receptor neurons (TRNs: ALML/R, AVM, PVM, and PLML/R) by using transgenic animals expressing channelrhodopsin (ChR2) under the *mec-4* promotor (Methods)^39^. In response to natural stimuli, the posterior TRNs (PVM and PLML/R) respond to posterior touch, inducing forward acceleration, whereas the anterior TRNs (ALML/R and AVM) respond to anterior touch, inducing reversals ^4,7,8,39,43^. To characterize the dynamics of these responses, we applied an m-sequence light stimulus to either the anterior or posterior region of transgenic animals, selectively stimulating the anterior or posterior TRNs, respectively (Fig 2A). We first computed linear filters characterizing the relationship between anterior TRNs and either discrete or continuous behavior (Fig 2 and Fig S1). As a control, we also performed experiments with animals that were not fed all trans-retinol (ATR), a cofactor required for ChR2 function. The computed filters for control animals are flat, zero-mean signals (Fig 2, gray lines). In contrast, the acceleration BWA for the ATR-fed worms results in a filter with a robust negative peak, −13 ± 0.50 μm/s^2^ (Fig 2Ei). The presence of this peak in the experimental group and its absence in the control group suggest that the filter is optogenetically induced, and not due to spontaneous behavior. We attribute small fluctuations as experimental noise rather than representing a true high frequency response. Lastly, this deceleration in the experimental group is expected from typical reversal responses to anterior touch stimulation^4,7^.

**Fig 2:**
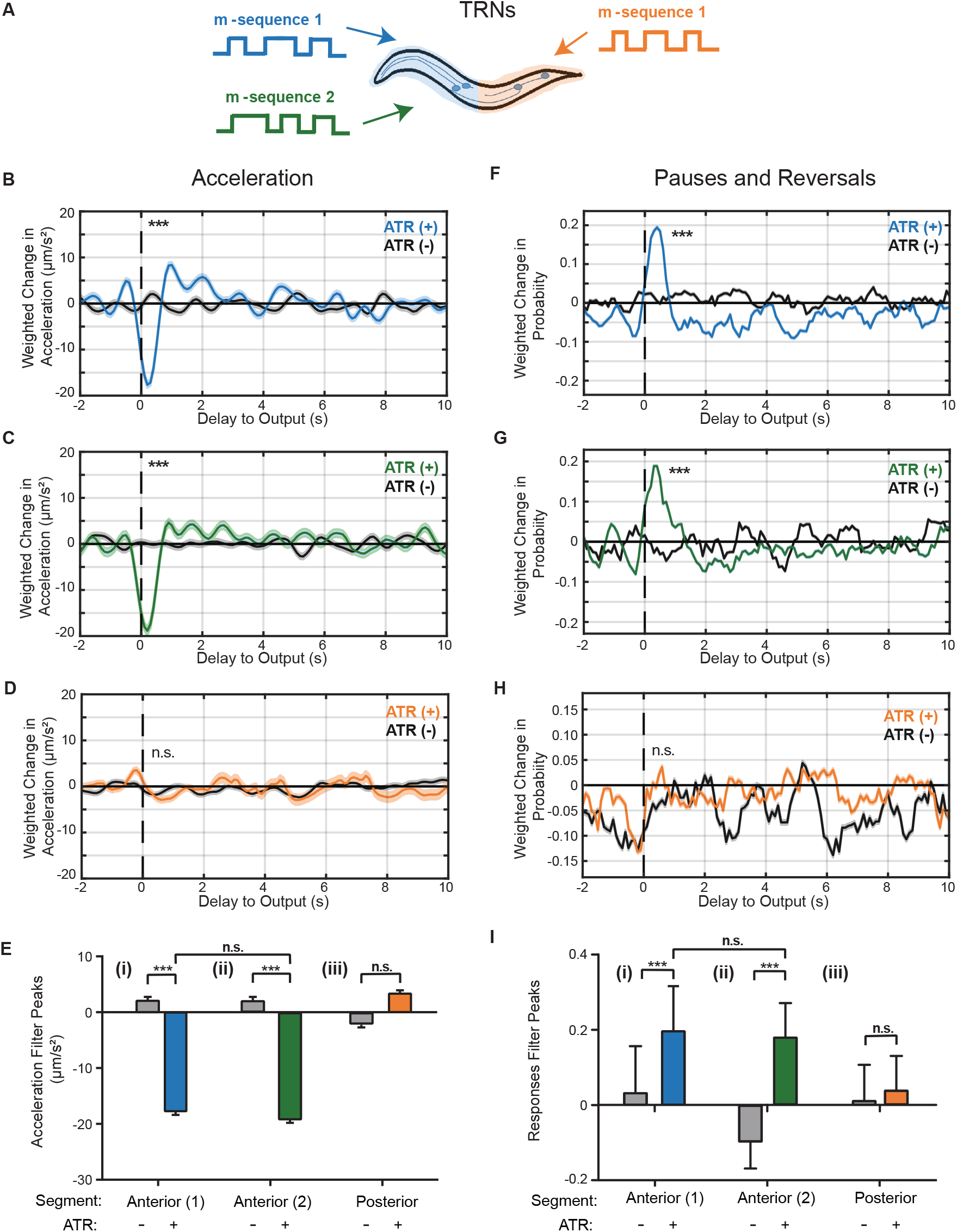
Linear filters for the touch receptor neurons (TRNs) responses are robust and reproducible. (A) Stimulus patterns and neurons being analyzed. Animals used in these experiments express channelrhodopsin using the mec-4 promoter (Methods). (B-D) Linear filters computed from BWA for acceleration when stimulating the anterior (B,C) or posterior (D) TRNs. (E) Comparisons of peak values from computed linear filters in B-D. (F-H) Linear filters computed from BWA for pauses and reversals when stimulating the anterior (F,G) and the posterior (H) TRNs with an m-sequence. (I) Comparisons of peak values from computed linear filters in F-H. Colored plots represent filters computed from ATR-fed animals, black plots represent filters computed from control (not ATR-fed) animals. Dark line and light shade represent BWA and SEM, respectively (sample sizes listed in Table S1). Error bars in bar plots indicate SEM (sample sizes listed in Table S1). Statistical significance for peaks computed using student’s t-test (***p<0.001) and statistical significance of filters computed using shuffled data (Fig S2).

In order to further assess the validity of the resulting filters, we performed statistical tests comparing true filters and filters computed from shuffled data (Methods). We compute magnitudes for all filters, defined as the L2 norm, to the correctly computed filter. Data is shuffled in four different ways (Methods). In all tests, the BWA computed from experimental data has the highest magnitude compared to filters computed from shuffled data (Fig S2). Together with the statistical comparison of ATR-fed and non ATR-fed animals, we conclude that the BWA for acceleration is robust and descriptive of the behavioral response.

In addition, our method also reveals new information about the dynamics of these responses. From the BWA, we can characterize metrics such as the delay to the peak (0.2s) and the decay timescale of the filter (0.4s); these temporal characteristics are critical for accurately predicting response to activation of the anterior TRNs. In comparison, the BWA computed with velocity also returns a linear filter with a negative peak (−6.1 ± 0.39 μm/s), although with a longer delay to peak (0.7s) and longer decay timescale (0.6s) (Fig S1). The difference between temporal characteristics of these two filters suggests that although animals reverse for a relatively long time after the stimulus (1.3s), the deceleration portion of this reversal only takes place in the first 0.6s after a stimulus.

To ensure that the computed linear filters are not an artifact from the input signal itself, we tested computing filters using a different m-sequence stimulus. Using acceleration as an example, we observe a similar linear filter to those obtained with the previous stimulus (Fig 2C, as compared to 2B). When comparing the peak values of the filters computed with different stimuli, there is no statistical difference (Fig 2Eii). These results demonstrate that the linear filters are indeed characteristic of *C. elegans’* behavioral output specifically in response to the activity in the anterior TRNs, and independent of the input signal.

Next, we sought to compare the dynamics of the animals’ response between anterior or posterior TRN activities. Previous findings have shown that applying a mechanical force to the posterior region of the animal induces an acceleration, and PLM is required for these responses ^4,7,8^. As with the anterior TRNs, we stimulated the posterior TRNs by applying an m-sequence light stimulus to the posterior half of the animal, and computed the BWA for the same quantified behaviors (Fig 2 and Fig S1). The filter for acceleration has a positive peak (2.8 ± 0.48 μm/s^2^), although with a much smaller magnitude than its anterior counterpart and is not statistically significant compared to non ATR-fed worms (Fig 2D). Additionally, the filter is not statistically significant when testing against filters for shuffled data (Fig S3). Interestingly, although the computed linear filter for the posterior TRNs has a peak in the direction that is consistent with previous findings, it is close to zero-mean. One interpretation that is consistent with literature is that worms have a lower rate of responses when activating PLM and PVM in comparison to the activating anterior TRNs. This is not surprising, as worms are generally moving forward and do not require a change in behavior to escape the weak stimulus, whereas avoidance of a weak anterior stimulus requires a directional change.

In addition to continuous signals, we also estimated linear filters for the probability of transitions between defined states. Unlike in predicting continuous variables (e.g. acceleration and velocity), filters computed for these behaviors indicate a change in probability of transitions to these behaviors. When computing the BWA with transitions into pauses or reversals in response to anterior TRNs, we observe linear filters with positive peaks that are statistically significant as compared to non ATR-fed animals (Fig 2F,G, I). Similarly, the filters computed from shuffled data support this statistical significance (Fig S3). This indicates that activating the TRNs induce an increase in probability of transitions to pauses or reversals, and this increased likelihood happens within the first second after a stimulus. In contrast, when stimulating the posterior TRNs, the filter computed for transitions into pauses and reversals is close to zero-mean, indicating that the stimulus does not alter these behaviors significantly (Fig 2H,I, and Fig S3).

### Reverse Correlation Analysis of Harsh-Touch Sensing PVD Neurons

In addition to the TRNs, *C. elegans* has another set of neurons that are responsible for body touch sensation. The PVD neurons are morphologically unique sensory neurons that have extensive and organized dendritic structures expanding most of the body of the worm; in contrast, TRNs are tiled (Fig 1A). Additionally, the PVD neurons are known to respond to harsh touch, as opposed to gentle touch or nose touch^5,10–13^. Because of the morphological and functional differences between the PVD and TRN systems, we ask whether there are also downstream differences in spatial and temporal behavioral response dynamics. To do so, we applied the same reverse correlation method to animals expressing ChR2 in the PVD neurons ^13^.

For comparison with the TRNs, we again divided the stimulus regions into anterior and posterior segments, and computed the BWA and estimated linear filters for the same behaviors (Fig 3A). Interestingly, when the animal is stimulated either anteriorly or posteriorly, the BWA’s for acceleration both have positive peaks (Fig 3B,C), indicating that activating either of these segments of PVD induces an increase in velocity. This positive peak is also observed for both segments when computing the BWA with velocity (Fig S4). However, only the filters from the posterior segment are statistically different from the non-ATR group, with a higher positive peak for both acceleration (Anterior 4.0 ± 0.58 μm/s^2^ vs Posterior 7.1 ± 0.58 μm/s^2^) and velocity (Anterior 3.5 ± 0.32 μm/s vs Posterior 7.1 ± 0.32 μm/s) (Fig 3D and Fig S4). When computing the BWA with transitions into pauses or reversals in response to either anterior or posterior PVD, we observe flat, zero-mean linear filters (Fig 3 E, F). These filters are statistically indistinguishable from the non-ATR fed control group (Fig 3G), indicating that activation of the PVDs do not induce a change in probability of these events. When comparing these filters with shuffled data, only the posterior acceleration filter is statistically significant (Fig S5). This contrast from the TRN filters suggests a different role for PVD sensory neurons in the behavioral circuit – that PVD activation promotes positive acceleration, and TRNs promote negative acceleration, consistent with previous findings ^8,12^.

**Fig 3:**
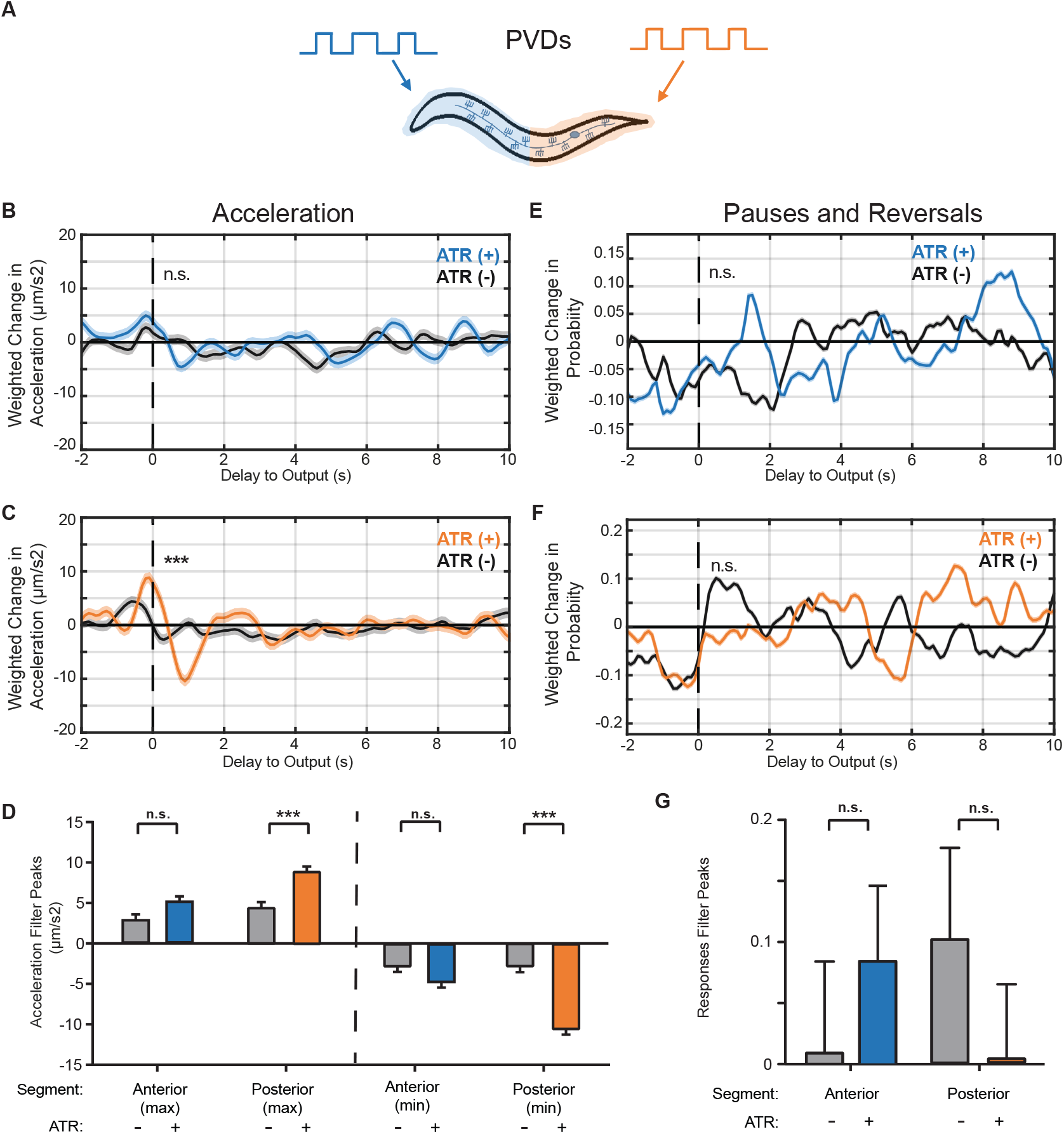
Linear filters for PVD activity illuminate dynamic differences between gentle and harsh touch systems. (A) Stimulus patterns and PVD neurons being analyzed. (B,C) Linear filters computed from BWA for acceleration when stimulating the anterior (B) or posterior (C) regions of PVD. (D) Comparisons of peak values from computed filters. (E,F) Linear filters computed for pauses and reversals when stimulating the anterior (E) and posterior (F) regions. (G) Comparisons of peak values from computed filters. Colored plots represent filters computed from ATR-fed animals, black plots represent filters computed from control (not ATR-fed) animals. Dark line and light shade represent BWA and SEM, respectively (sample sizes listed in Table S1). Error bars in bar plots indicate SEM (sample sizes listed in Table S1). Statistical significance for peaks computed using student’s t-test (***p<0.001) and statistical significance of filters computed using shuffled data (Fig S4).

In addition to the magnitudes, the context of peak occurrence can also be informative. The PVD acceleration filters have significant negative peaks following the positive peaks; the magnitudes of the negative peaks are of similar values to the first positive peak (Fig 3D). This suggests that the acceleration in response to PVD activation is more likely to occur when preceded by a negative acceleration. In other words, worms that are slowing down or reversing are more likely to respond to PVD activation and produce a positive acceleration. In contrast, the anterior TRN acceleration filters only contain one significant peak. These differences in the acceleration filters further supports the idea that PVD and TRNs influence different aspects of behavior.

### Linear-Nonlinear Models Predict Behavioral Response

In general, the filters computed from BWA in response to a white noise signal capture the linear dynamics of the analyzed systems. However, biological systems are rarely linear^24^. A common approach for modeling the nonlinear dynamics of a system is to use a linear-nonlinear cascade, where a static nonlinear filter is used to characterize the nonlinear dynamics not captured by reverse correlation ^20,23–25^. To define static nonlinear filters, we used the linear filters computed from BWA and compared predicted outputs with measured experimental outputs (Methods). For instance, for acceleration in response to anterior TRNs, we first compared predicted values with the quantified experimental values (Fig 4A, gray circles). Not surprisingly, there is a positive correlation between predicted and experimental outputs, indicating that the model does indeed capture linear dynamics in these responses. To capture the nonlinear dynamics of the response, we fit a static filter using a simple quadratic function (Fig 4A blue lines, Methods). Similarly, we also characterized nonlinear filters for velocity (Fig S6A) and transitions into pauses or reversals (Fig 4B, Methods). The quadratic functions greatly improve the model fit to the data, suggesting that they capture a large portion of the nonlinear dynamics of the anterior TRNs. We also computed static nonlinear filters for stimulation of the posterior TRNs. In comparison to the anterior TRNs, there is a lower correlation between experimental measurements and predicted values (Fig 4C,D, Fig S6B). This is expected, as the estimated linear filters for these neurons were close to zero-mean, yielding a small range of predicted responses. Furthermore, because the linear filters alone led to a low predictability of responses for posterior TRNs, nonlinear functions also fail to capture a large portion of the variability in responses (Fig 4C,D, Fig S6B, orange lines).

**Fig 4:**
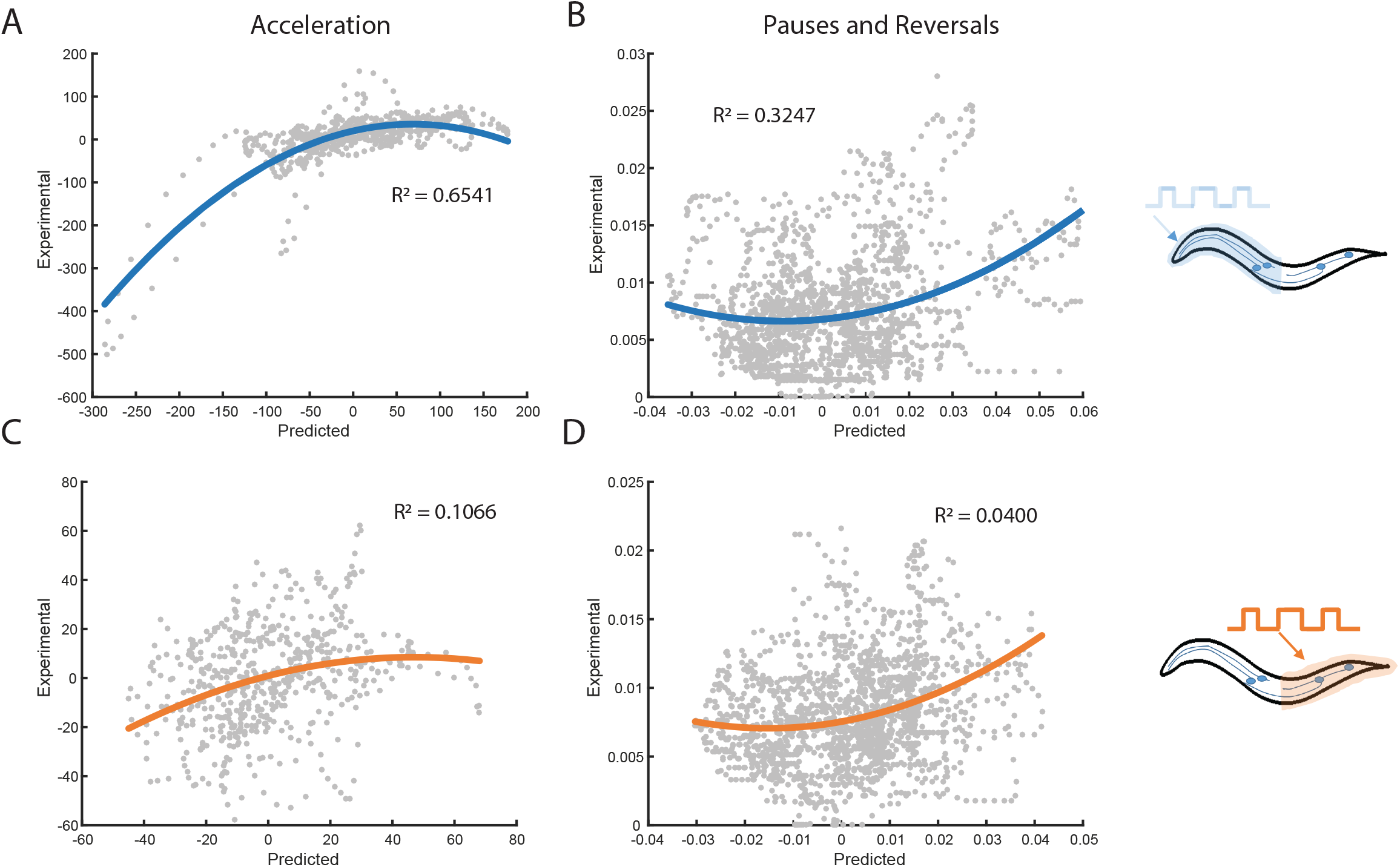
Static nonlinear filters capture nonlinear dynamics in behavioral outputs. Estimation of static filters to capture nonlinear dynamics. (A,B) Static nonlinear filters fitted using predicted values from the linear filter (x-axis) and experimental values (y-axis) when stimulating the anterior TRNs, for (A) acceleration and (B) transitions into pauses and reversals. (C,D) Static nonlinear filters when stimulating the posterior TRNs, for (C) acceleration and (D) transitions into pauses and reversals. Linear filters and experimental values are subsets of data used in Figure 2 (n=600 for all conditions). Colored traces represent computed nonlinear filters and gray dots represent independent time-points from measured and predicted values. Probability of discrete events is computed as the probability of an event occurring at a given time point.

We next sought to test the validity of using linear-nonlinear (LN) cascade models to predict behavioral responses to novel stimuli. To do this, we probed the anterior TRNs with a different m-sequence stimulus from the one used to compute the filters (Fig 5A, Methods). We first compared the measured velocities of animals to the predicted velocities when using the linear filter only (Fig 5B). Although the magnitude of predicted velocity from the model did not exactly match the experimental measurements, the model captures large features of the temporal dynamics of velocity in response to this novel stimulus. Next, we incorporated the static nonlinear filter to predict velocities (Fig 5C). When using the LN model, the magnitudes of predicted velocities are more similar to experimental values, leading to more accurate predictions. In addition to predicting the continuous velocity of the animals, we also tested L and LN models for pauses and reversals, and observe predicted increases in probability of events similar to experimental values (Fig S7A,B). Incorporating the nonlinear component to these models also improves the model predictability.

**Fig 5:**
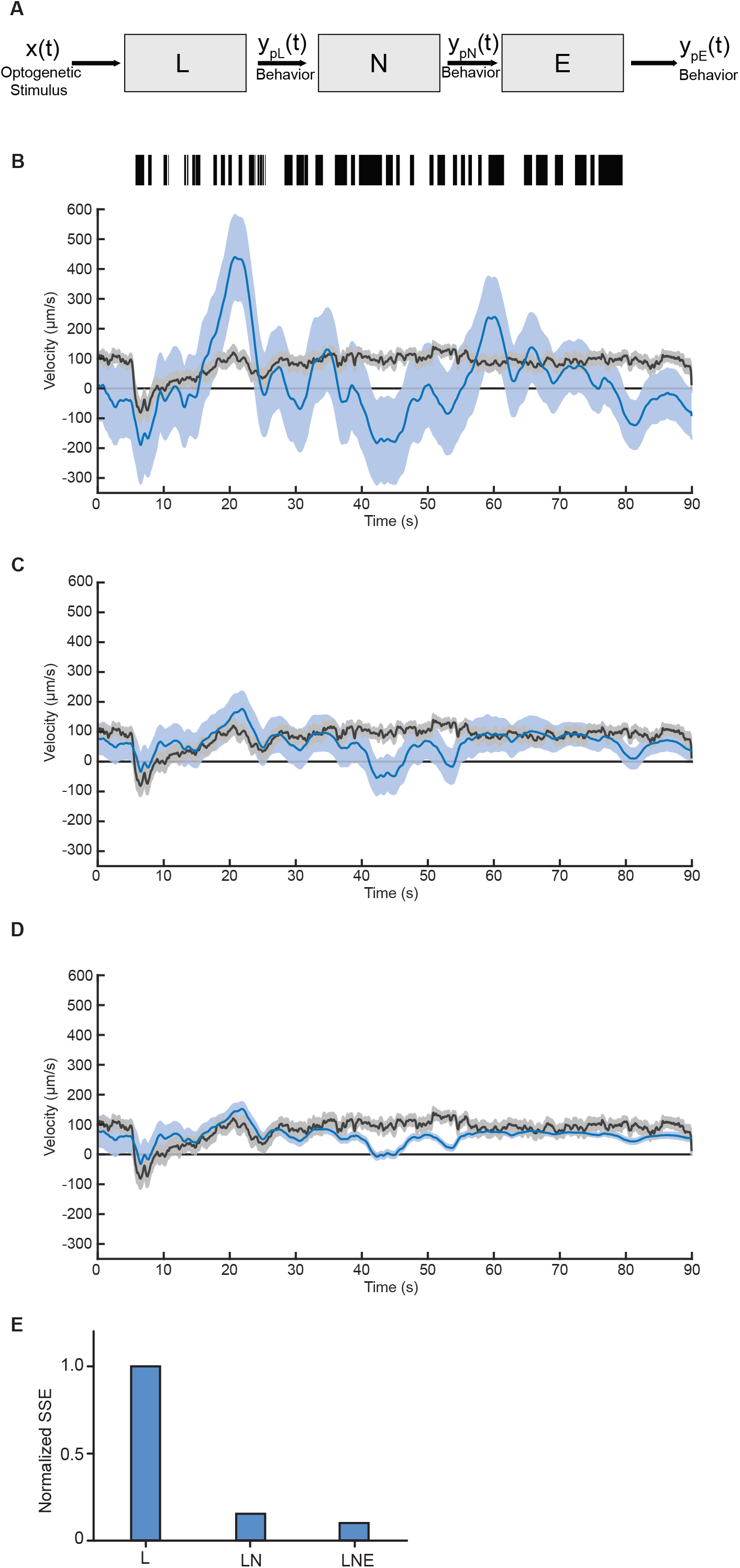
Linear-Nonlinear-Exponential (LNE) model accurately predicts behavioral response. (A) Block diagram of LNE model for used to predict behavioral responses to mechanosensory neuron activity: a LTI system modeled from BWA, followed by a static nonlinear filter and exponential decay filter. (B-D) Predictions of velocity for L (B), LN (C), and LNE (D) models (blue) and experimental traces (black). For experimental data, dark line and shade represent average and SEM, respectively (n = 31 animals). For model predictions, dark line represents model prediction and shaded area represents the 95% confidence interval (Methods). (E) Comparison of performance of models, computed as the sum of squared error (SSE) and normalized to the linear model performance value (Methods).

Interestingly, in our experiments we observe a time-dependent decrease in the magnitude of responses, which fails to be captured in time-scales of the dynamic linear filters (Fig 5B,C latter half). Biologically, this habituation of responses is commonly observed in sensory systems ^44^. In general, although LN models can predict system responses, this is true only to the time-scales captured in the linear filters, and does not capture adaptation dynamics. To model this decay of responses, we add a dynamic exponential function following the LN cascade (Fig 5A). We tested a wide range of decay rate values using this model and found that a decay rate of 50s best provided the best predictions (Fig S8). Interestingly, this decay rate is consistent with previous findings from investigations of habituation to stimulation of TRNs, with both tapping and optogenetic stimuli ^45^. When adding this exponential component to our model, the accuracy of our model’s predicted behavioral responses improves for later time points of the trials, thus improving the overall accuracy of our models (Fig 5D, Fig S7C,D). These results illustrate how the linear filters computed from BWA, when combined with additional nonlinear filters, are powerful in predicting temporal dynamics of behavioral responses to sensory neuron activation, and likely generalizable to other sensory responses.

### Spatially Refined Selective Illumination Improves Resolution of Linear Filters from BWA

We have thus far characterized mechanosensory systems by probing either the anterior or posterior segments of the animal, similar to previous investigations of the receptive fields of mechanosensory systems ^12,14^. To further examine the spatial resolution of the mechanosensory systems, we took advantage of our selective-illumination light stimulus, which allows for the probing of specific spatial segments as small as 14μm ^39^. We characterized the TRN system with better resolution by increasing the number of segments in our stimulus to 4 (Fig 6A). We applied an m-sequence stimulus selectively to one of the four segments, and computed linear filters for both continuous and discrete behavioral outputs (Fig 6 and Fig S9). This particular discretization of the TRN system allows for the computation of separate filters for the processes and cell bodies of ALM and AVM, as well as separate filters for PVM and PLM cell bodies (while keeping a high number of photons in the stimulus region). We first computed filters for acceleration in response to stimulating four segments. The filters for the most anterior quarter and second-most anterior quarter have a prominent negative peak, statistically significant when compared to non-ATR fed animals (Fig 6B,C,F). These filters are also statistically significant when compared to shuffled data (Fig S10). Interestingly, these filters are similar to the filter computed from stimulating the entire anterior region (compare to Fig 2B,C). This suggests that there are no observable differences in acceleration dynamics between cell body and axon activity of the anterior TRNs.

**Fig 6:**
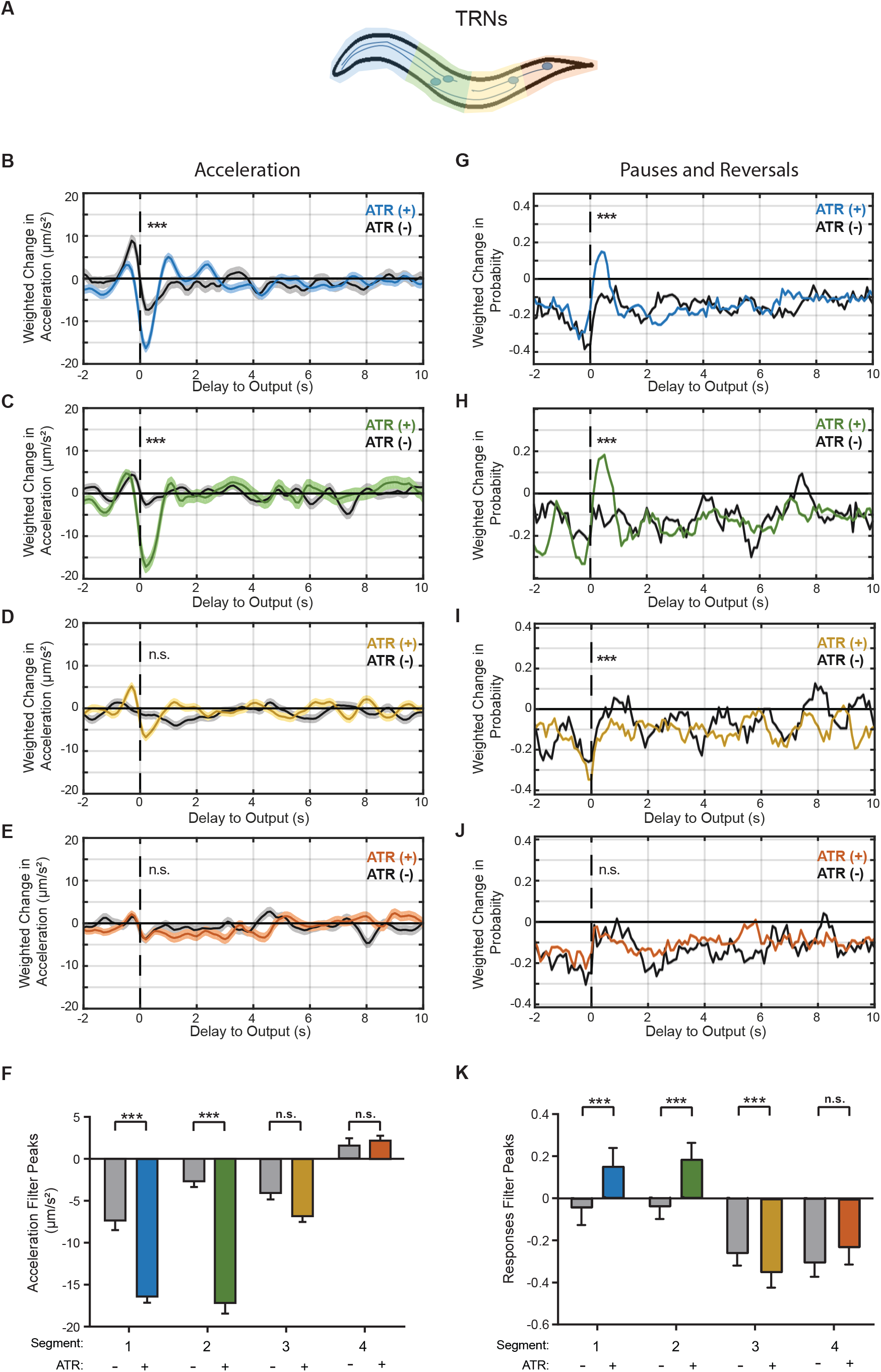
Decreasing the size of stimulus region allows for the estimation of a spatiotemporal receptive field with higher resolution. (A) Stimulus patterns used to analyze TRNs with improved spatial resolution. (B-E) Linear filters computed for acceleration when stimulating the most anterior (B), the second-most anterior quarter (C), second-most posterior quarter (D), and the most posterior quarter (E) of the TRNs with an m-sequence. (F) Comparisons of peak values from computed filters in B-E. Error bars indicate SEM (sample sizes listed in Table S1). (G-J) Linear filters computed for acceleration when stimulating the most anterior (G), the second-most anterior quarter (H), second-most posterior quarter (I), and the most posterior quarter (J) of the TRNs with an m-sequence. Colored plots represent filters computed from ATR-fed animals, black plots represent filters computed from control (not ATR-fed) animals. Dark line and light shade represent BWA and SEM, respectively (sample sizes listed in Table S1). (K) Comparisons of peak values from computed filters in B-E. Error bars indicate Standard deviation. Statistical significance for peaks computed using student’s t-test (***p<0.001) and statistical significance of filters computed using shuffled data (Fig S10).

Not surprisingly, the filters for acceleration in response to the most posterior quarter and second most posterior quarter are both flat and are statistically indistinguishable from filters computed with non-ATR fed animals (Fig 6D,E,F). These filters are also not statistically significant when comparing to shuffled data (Fig S10). Similar to the anterior region, the acceleration filters for the separate posterior segments are similar to the flat filter computed from stimulating the entire posterior region (compare to Fig 2D).

We next computed linear filters for transitions into pauses or reversals, and found differences in spatial encoding. The results for the anterior segments did not reveal much spatial encoding, with the filters for both the most anterior quarter and second-most anterior quarter both having positive peaks (Fig 6G,H,K), similar to the filter computed when stimulating the entire anterior regions (compare to Fig 2F,H). These filters are also statistically significant when comparing to shuffled data (Fig S10). This suggests that there is low spatial encoding of these discrete behavioral responses between the axons and cell bodies of the anterior TRNs. Interestingly, we observe different filters when dividing the posterior segment of the TRNs into separate segments for the cell bodies of PVM and PLM. The filter for the most posterior quarter, which includes the PLM cell body, is again a flat filter (Fig 6J), similar to the filter computed when stimulating the entire posterior region (compare to 2H). Surprisingly, the filter for second-most posterior quarter has a negative peak, statistically significant when compared to non-ATR fed animals (Fig 6I,K). This filter is also statistically significant when compared to shuffled data (Fig S10). The negative peak indicates that there is a reduced probability of pauses and reversals when activating PVM cell body. This suggests that PVM potentially has a previously undescribed function of inhibiting pauses and reversals. Additionally, the difference in filters for the four segments implies that the TRNs employ their tiled network to allow for spatial encoding of behavioral responses. This suggests that the morphological differences between the tiled TRNs and branched PVDs are used to differently control downstream activity.

## Discussion

The nervous system continuously transduces sensory stimuli into neuronal activity and appropriate behavioral outputs. One of the biggest challenges in mapping this neuronal encoding is the lack of a quantitative framework for characterizing how a layer of neural activity is transduced into the downstream circuit. In this work, inspired by previous work in modeling neuronal systems, we built a framework that uses reverse correlation analysis with a custom tracking platform to analyze a *C. elegans* sensory system. We investigated the spatial and temporal encoding of two mechanosensory systems, the gentle touch sensing TRNs and the harsh touch sensing PVDs. We computed several linear filters that quantitatively describe transformations between sensory neuron activity and behavioral outputs, and support previous findings about the systems. Analysis of the PVDs produced linear filters that indicate an increase in velocity and acceleration from their activation, which is consistent with literature on its function ^5,10–13^. Similarly, the linear filters computed for the TRNs were also consistent with previous literature: the anterior TRNs show decreases in velocity and acceleration, and an increase in probability of pauses and reversals ^4,7,8,39,43^, and the posterior TRNs show an increase in acceleration ^4,8,39^. It should be noted that we do not measure expression levels of ChR2 in the sensory systems, and any differences in computed filters could be explained by differences in expression levels. However, when assuming uniform expression levels across the sensory systems, our results provide spatiotemporal receptive fields for these systems that are consistent with previous findings ^7^.

The linear filters resulting from our method provide several insights into the circuitry and morphological differences between the two sensory systems. First, although we used identical stimuli for both segments, the filters produced from activating the anterior TRNs were much more robust than the filters from activating the posterior TRNs, suggesting that downstream interneurons in this circuit are more responsive to the anterior neurons. This preference in downstream activity has also been observed in experiments involving tap responses, which show that reversals dominate over accelerations when tapping cultured plates, and this preference occurs downstream of sensory neuron activity ^8^. In contrast, the filters for the posterior segments of PVD were more robust than the anterior segments. This is also consistent with previous findings that show PVD is required for posterior harsh touch sensation, but not required for anterior harsh touch ^12^. A key difference in our experiments is that we bypass mechanoreceptor activation, and can therefore separate out effects due to differences in sensory neuron response to different spatial stimuli, as well as other neurons that might affect response rate. Therefore, one possible mechanism for the differential decision-making is that the two mechanosensory systems may have different strengths of connections to postsynaptic command interneurons. Particularly for PVD, although the number of physical synapses to forward command neuron PVC and backward command neuron AVA are similar ^46^, the functional connectivity seems to be higher for PVC compared to AVA. Our results strongly support this hypothesis.

Our results also provide insight on the levels of spatial encoding in the TRNs and PVD systems. The TRNs, which employ a tiled network to cover the body, appear to have more spatial encoding. When comparing the computed filters for the anterior and posterior TRNs, most behaviors show distinct differences in responses. Furthermore, when analyzing this system in four segments, we observed differences in linear filters among the four segments. In contrast, the branched network in PVD does not appear to spatially encode behavioral responses. The filters from activating the anterior and posterior segments of the PVD system have similar dynamics, with the anterior filters having slightly smaller magnitudes and longer delays. This contrast between the two mechanosensory systems suggests that although both the TRNs and PVDs have spatially distributed processes to sense touch throughout the body, the unique morphological strategies in the two systems lead to differences in their capabilities of encoding responses. Biologically, this disparity in encoding can be explained by their morphologies and perhaps synaptic connectivity to downstream neurons, as the tiled TRN system consists of more nodes, which could allow for more spatially specific behavioral responses.

One new finding from our experiments concerns the role of the cryptic PVM neuron. Although shown to respond to mechanical stimuli ^47^, its role in mediating behavior is poorly understood ^4,7,8,39^. We found that activating PVM did not induce significant changes in velocity, but induced a slight decrease in acceleration. Interestingly, PVM activation significantly reduced the probability of reversal events. These filters suggest a unique function for PVM in modulating escape response. In contrast to the other TRNs, PVM does not induce escape responses, but rather suppresses these behaviors, as well as decrease the velocity of forward movement.

The findings in this work demonstrate the utility of our method for providing new insights into the dynamics of the mechanosensory system in *C. elegans*, one of the earliest and better characterized neural circuits. By using a quantitative framework to compare the dynamics between the two sensory systems, we recapitulated qualitative findings from previous literature, and provide further insights in the temporal and spatial encoding in these systems. Additionally, we used linear filters computed from BWA to create LNE models that can predict the behavioral responses of animals in response to activity in sensory neurons alone. Because this method is noninvasive and independent of natural stimulus, it can be easily extended to investigate the dynamics of other neural circuits in *C. elegans* and other model organisms. We foresee many potential applications in better understanding sensory behavior responses and sensory integration.

## Methods and Materials

### *C elegans* Culture and Maintenance

We used transgenic worms expressing channelrhodopsin in various mechanosensory neurons. Worm populations were cultured at 20C in the dark on standard nematode growth medium (NGM) petri dishes. Plates were coated with OP50 bacteria lawn supplemented with the cofactor required for channelrhodopsin, all-*trans*-retinal (Sigma-Aldrich). The solution was prepared by diluting a 50mM stock solution (in ethanol) in OP50 suspension to a final concentration of 100uM. Control animals were grown in parallel on OP50 without all-*trans*-retinal. All worms tested were F1 progeny of P0 adults picked onto seeded plates 3-4 days before experiments. Animals were washed to unseeded NGM plates 1hr prior to assays. Animals were then picked to individual plates for experiments. Each animal was exposed to a single stimulus profile and then discarded. The strains used in this work included AQ2334: *lite-1(ce314); ljls123[pmec-4::ChR2; punc-122::rfp]* ^39^ and ZX899: *lite-1(ce314); ljls123[pmec-4::ChR2; punc-122::rfp]* ^13^.

### Tracking and Light Delivery Platform

Experiments were performed on a tracking system adapted from a previously developed projector based microscopy system ^39^. The system uses an inverted microscope (Leica-DMIRB) with a low-magnification objective (x4) to image freely moving animals. We image using near-infrared light by applying a long-pass filter (715nm) to the transmitted light path and capture images using a large sensor NIR camera (Basler acA2040-180kmNIR), which limits interference in blue light used for optogenetics stimulus. A three-color LCD projector is used as the light source for optogenetic stimulus with selective illumination. We use a camera with large sensor area to capture the full body of the animal, and use a small ROI and binning to reduce the size of images to improve processing speed and therefore tracking rate. A Lenovo desktop computer with an Intel Core i74790 Processor (8MB Cache, up to 4.0GHz) and a 512GB Solid State Drive and 16GB RAM was used to process images for tracking and selective illumination. Tracking of individual animals was performed by using images taken with the camera, and processed to compute the centroid of the animal in terms of x-y pixels on the camera FOV. Based on the position of the computed centroid, a command is sent to a motorized stage to move the animal to the center of the FOV. To apply a light stimulus with spatial and temporal control, we used a modified projector as the light source to the microscope. Images taken with the camera are processed to determine the outline of the animal’s body in each frame. The appropriate illumination pattern is then computed and sent to the projector. Stimuli were only presented when ansterior and posterior segments were correctly computed by the algorithm; during pirouettes or other uncommon postures, stimulus presentation was paused. This process was performed at a rate of 13 frames per second. For each animal, illumination profile and tracking videos were saved for future analysis.

### Quantitative Behavior Analysis

To extract quantitative behavioral features from tracking recordings, a custom MATLAB script was developed. A series of segmentation and morphological processes were used to extract body postures in each frame. We combined extracted postures with recorded stage movements to quantify several behaviors. We computed various “continuous” behaviors that have a scalar value for each time point. This includes velocity (magnitude), velocity (angle), acceleration, head angle, angular velocity. We also classified various “discrete” behaviors that have been used in previous works ^18,40,48,49^. These include behaviors such as pauses, reversals, omega turns, and turns. Each of these behaviors were classified by applying thresholds on quantified continuous behaviors. Pauses and reversals were classified by applying both vertical and horizontal thresholds on velocity measurements. Omega turns were classified by applying a threshold on the eccentricity of the animal’s posture. Curves were classified by applying a threshold on the angle of position trajectory.

### White Noise Experiments

We used the selective illumination capability of the tracking system to deliver spatially controlled light stimuli to freely moving animals expressing ChR2 in their mechanosensory neurons. We used a pseudorandom m-sequence, a binary signal with unbiased spectrum, with similar properties to a Gaussian white noise signal ^22,31^. We tested several white noise signals, and found that an m-sequence with a maximum frequency of 2Hz produced reliable results, as it allows for testing time scales appropriate for behavioral responses. We use a light intensity of 0.75mW/mm^2^ as it induces reliable and varying behavioral responses, similar to previous work ^39^. The generated pseudorandom sequences were repeats of a 6-bit words, 63 value length m-sequences (2*(2^6^-1) = 126 values). We deliver the same pseudorandom signal for each experimental group, applying the signal through the tracking system and changing values in the m-sequence at 2Hz, which is lower than the Nyquist Frequency (acquisition rate is 13Hz). Stimuli were only presented when ansterior and posterior segments were correctly computed by the tracking algorithm; during pirouettes or other uncommon postures, stimulus presentation was paused.

### Reverse Correlation Analysis

To compute mathematical functions that describe the transformations from sensory neuronal activity into behavior, we first modeled the entire animal as a linear transducer:

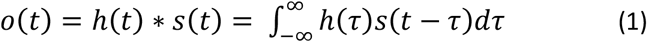

where the relationship between the input signal (neuronal activity through optogenetics) s(t) and output signal (behavior) o(t) is characterized by a function h(t). We assume that the system is causal, and h(t)<0 for t<0. We used standard reverse-correlation similar to ^29–31,34,35^, and computed h(t) for specific behaviors by computing a “behavior-weighted-average” (BWA):

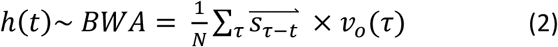

where the stimulus preceding each time-point is weighed by the scalar value of the behavior at that time. We convert the light stimulus patterns into −1 and 1 for when the light is on and off, respectively. For continuous behaviors, we used the scalar values at each time points as the weights. For discrete behaviors, we used a binary signal indicating transitions from forward movement to specific states. For all cases, we compute linear filters using 400 points preceding and following each time point (801 total timepoints). The points preceding each time point are computed as a control to capture experimental variablity.

### Statistical Significance of Computed Filters

Behavior-weighted averages (BWAs) were tested for significance by comparing their magnitude, computed as the L2 norm, to a distribution of random filters computed with shuffled data. We tested four different methods of shuffling data: cyclic shuffling of the stimulus vector by a random integer, cyclic shuffling of the output vector by a random integer, random permutations of the stimulus vector, and random permutations of the output vector. For each test, we perform the same computation with the shuffled data and repeat 100 times. The BWA is classified as significant if its magnitude is higher than all shuffled data tests. Random integers were generated from a uniform distribution from 1 to length of vector using the MATLAB function rand, and random permutations of vectors were performed using the MATLAB function randperm.

### Nonlinear Filters and Model Predictions

We model static nonlinear filters for each behavioral response in order to extract the nonlinear dynamics not captured in the linear filters computed from reverse correlation ^50^. We first compute linear model predictions by convolving the computed linear filters from presented stimuli in each trial used, as shown in equation (1). We then compare these linear predictions to the measured outputs at each time point, and fit a quadratic function. For “discrete” behaviors, probabilities for transitions into specific behaviors were calculated at each time point. These quadratic functions are then used as static nonlinear filters in a linear-nonlinear (LN) cascade model for specific behavior transformations.

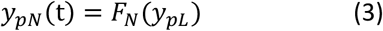

where the predicted nonlinear output is a static function of the predicted linear output. We also apply an exponential decay filter (LNE) to capture nonlinear adaptations to the stimuli. We apply this exponential factor to only the changes in behavior after the stimulus:

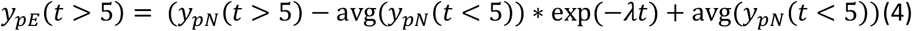

where the decay parameter *λ* is 50s, based on empirical data (Fig S8) and previous findings^45^. We use bootstrap sampling to compute 95% confidence intervals for our model predictions. Confidence intervals were computed using the MATLAB functions bootstrp and bootci, computed with 1000 resamples of the stimulus data.

### Statistics

Linear filters are presented as mean ± SEM as computed by the BWA. The two-tailed Student’s t-test was used to compare filter peaks between two groups. Peaks were determined by searching for local maxima in the filters between −1 < t < 1. P<0.005 was considered statistically significant. Accuracy of best-fit nonlinear filters were computed as coefficients of determination (R^2^ values). Performance of models were compared using the sum of squared error (SSE). Values are normalized to the SSE value for linear models.

### Code Availability

All custom code used to generate results in this manuscript are available on Github (https://github.gatech.edu/pages/dporto3/BWA-v1/).

### Data Availability

All behavior and stimulus data generated during the current study are available from the corresponding author upon reasonable request.

## Supporting information

## Acknowledgments

The authors gratefully acknowledge the funding support of the US National Institutes of Health (R01NS096581, R01GM088333, R21EB021676, and R21EB020424 to HL).

## Author Contributions

DP and HL designed experiments. DP, JG, and YZ performed experiments. DP wrote the manuscript and prepared all figures. All authors reviewed the manuscript.

## Supplementary Information Legends

**Movie S1:** Example trial of white noise stimulation in our platform. An m-sequence light signal is delivered to either the anterior or posterior segment of the animal while simultaneously being tracked. For each trial, various discrete and continuous behaviors are quantified (Methods).

**Movie S2:** Sample filter computed using BWA as a function of sample size used for the computation.

**Figure S1: Additional linear filters for TRNs.** Linear filters computed for various behaviors when stimulating the anterior TRNs with an m-sequence signal (left), a different m-sequence signal (center), and the posterior TRNs (right). Dark line and light shade represent BWA and SEM, respectively. Colored plots represent filters computed from ATR-fed animals, black plots represent filters computed from control (not ATR-fed) animals. Sample sizes listed in Table S1.

**Figure S2: Comparison of shuffled data significance tests.** Results from comparison of four methods of shuffling data for statistical significance tests of linear filters. (A) Cyclic shuffling of stimulus vector by a random integer. (B) Cyclic shuffling of behavior vector by a random integer. (C) Random permutation of stimulus vector. (D) Random permutation of behavior vector. Bar plots represent the magnitude of filters, computed as the L2 norm, and are plotted in ranked order from highest to lowest magnitude. Colored bar represents appropriately computed filter, gray bars represent filters computers with shuffled data.

**Figure S3: Significance test results for linear filters for TRNs.** Results from shuffled data significance tests for linear filters computed for activation of TRNs in Figure 2. (A-C) Significance test results for computed filters for acceleration for anterior TRNs (A,B) and posterior TRNs (C). (D-F) Significance test results for computed filters for pauses and reversals for anterior TRNs (D,E) and posterior TRNs (F). Bar plots represent the magnitude of filters, computed as the L2 norm, and are plotted in ranked order from highest to lowest magnitude. Colored bar represents appropriately computed filter, gray bars represent filters computers with shuffled data.

**Figure S4: Additional linear filters for PVDs.** Linear filters computed for various behaviors when stimulating the anterior (left) and posterior (right) PVDs with an m-sequence signal. Dark line and light shade represent BWA and SEM, respectively. Colored plots represent filters computed from ATR-fed animals, black plots represent filters computed from control (not ATR-fed) animals. Sample sizes are listed Table S1.

**Figure S5: Significance test results for linear filters for PVD.** Results from shuffled data significance tests for linear filters computed for activation of PVD in Figure 3. (A,B) Significance test results for computed filters for acceleration for anterior (A) and posterior (B) segments of PVD. (C,D) Significance test results for computed filters for pauses and reversals for anterior (C) and posterior (D) segments of PVD. Bar plots represent the magnitude of filters, computed as the L2 norm, and are plotted in ranked order from highest to lowest magnitude. Colored bar represents appropriately computed filter, gray bars represent filters computers with shuffled data.

**Figure S6: Static nonlinear filters for velocity.** Static nonlinear filters fitted for predicted values from the linear filter (x-axis) against experimental values (y-axis) when stimulating the anterior TRNs. Linear filters and experimental values are subsets of data used in Figure 2 (n>1,730 for all conditions). Colored traces represent computed nonlinear filters and gray dots represent independent time-points from measured and predicted values.

**Figure S7: Model predictions of reversal initiations.** Comparison of model predictions of reversal transitions (blue) and experimental traces (black) when using A) only the linear filter, B) a linear-nonlinear (LN) model, and C) an additional exponential component (LNE). For experimental data, dark line and shade represent average and SEM, respectively (n = 31 animals). For model predictions, dark line represents model prediction and shaded area represents the 95% confidence interval (Methods). Probability of reversal transitions is computed as the average of animals initiating a reversal at that time point. D) Comparison of performance of models, computed as the sum of squared error (SSE) and normalized to the linear model performance value.

**Figure S8: Comparison of decay factors.** Comparison of model predictions of velocity for various exponential decay factors. Exponential decays of 2.5s, 5s, 50s, and 100s were tested, with 50s showing the best fit. Performance of models is computed as the sum of squared error (SSE), normalized to the linear model performance value.

**Figure S9: Additional filters for spatially refined analysis of TRNs** Linear filters computed for various behaviors when stimulating the most anterior quarter (left), the second-most anterior quarter (second from left), the second-most posterior quarter (second from right), and the most posterior quarter (right) of the TRNs with an m-sequence signal. Dark line and light shade represent BWA and SEM, respectively. Colored plots represent filters computed from ATR-fed animals, black plots represent filters computed from control (not ATR-fed) animals. Sample Sizes are listed Table S1.

**Figure S10: Significance test results for linear filters for refined TRN analysis.** Results from shuffled data significance tests for linear filters computed for activation of TRNs in Figure 6. (A-D) Significance test results for computed filters for acceleration for most anterior (A), second-most anterior (B), second-most posterior (C), and most posterior (D) segments of TRNs. (E-H) Significance test results for computed filters for pauses and reversals for most anterior (E), second-most anterior (F), second-most posterior (G), and most posterior (H) segments of TRNs. Bar plots represent the magnitude of filters, computed as the L2 norm, and are plotted in ranked order from highest to lowest magnitude. Colored bar represents appropriately computed filter, gray bars represent filters computers with shuffled data.

**Table S1:** Sample sizes for computed linear filters in figures 2, 3, 4, 6, S1, S2, and S9.

