## Supplementary material for "Reverse-Correlation Analysis of the Mechanosensation Circuit and Behavior in *C. elegans* Reveals Temporal and Spatial Encoding"

Movie S1: Example trial of white noise stimulation in our platform. An m-sequence light signal is delivered to either the anterior or posterior segment of the animal while simultaneously being tracked. For each trial, various discrete and continuous behaviors are quantified (Methods).

Movie S2: Sample filter computed using BWA as a function of sample size used for the computation.

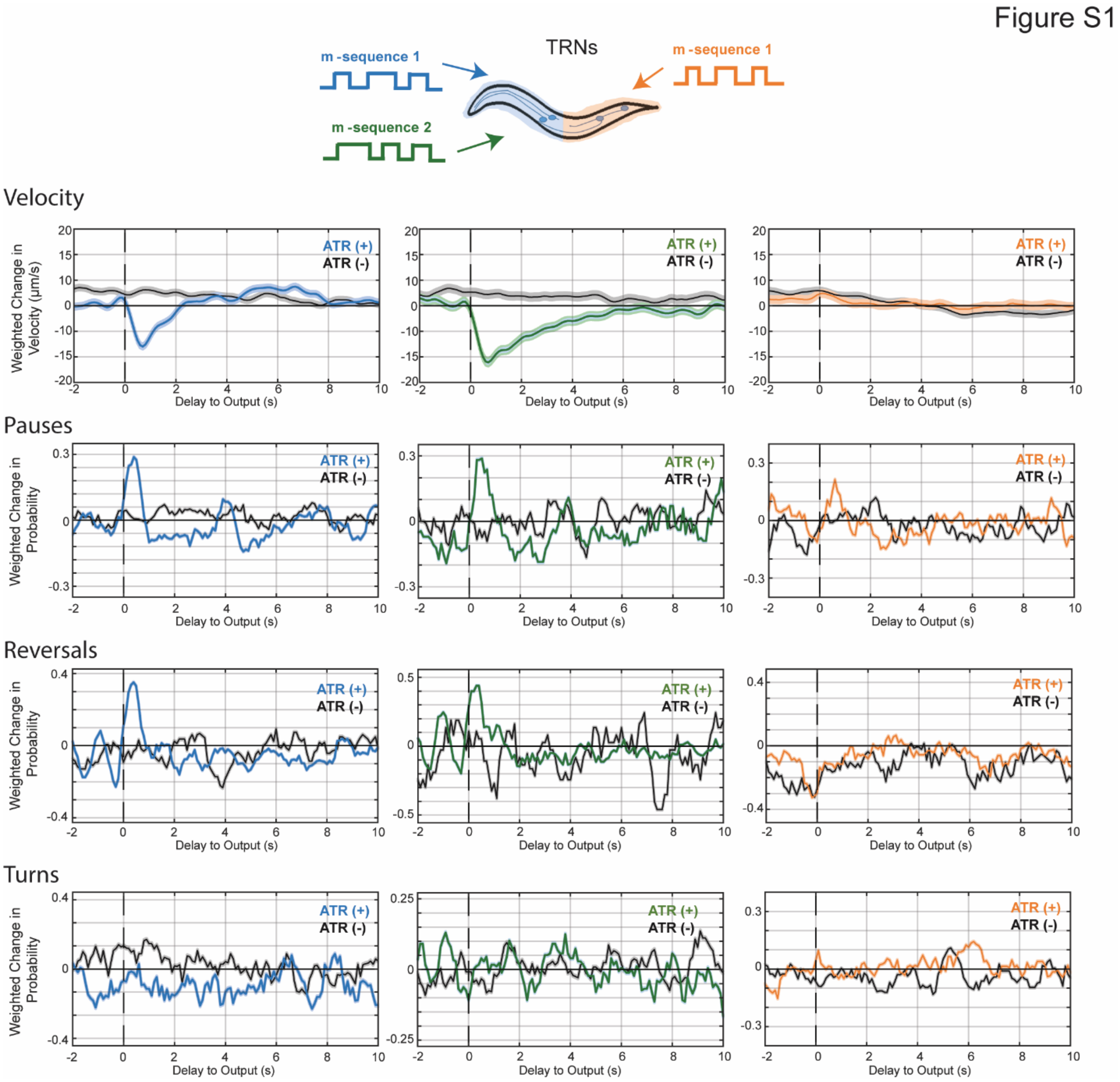


Figure S1: Additional linear filters for TRNs. Linear filters computed for various behaviors when stimulating the anterior TRNs with an m-sequence signal (left), a different m-sequence signal (center), and the posterior TRNs (right). Dark line and light shade represent BWA and SEM, respectively. Colored plots represent filters computed from ATR-fed animals, black plots represent filters computed from control (not ATR-fed) animals. Sample sizes listed in Table S1.

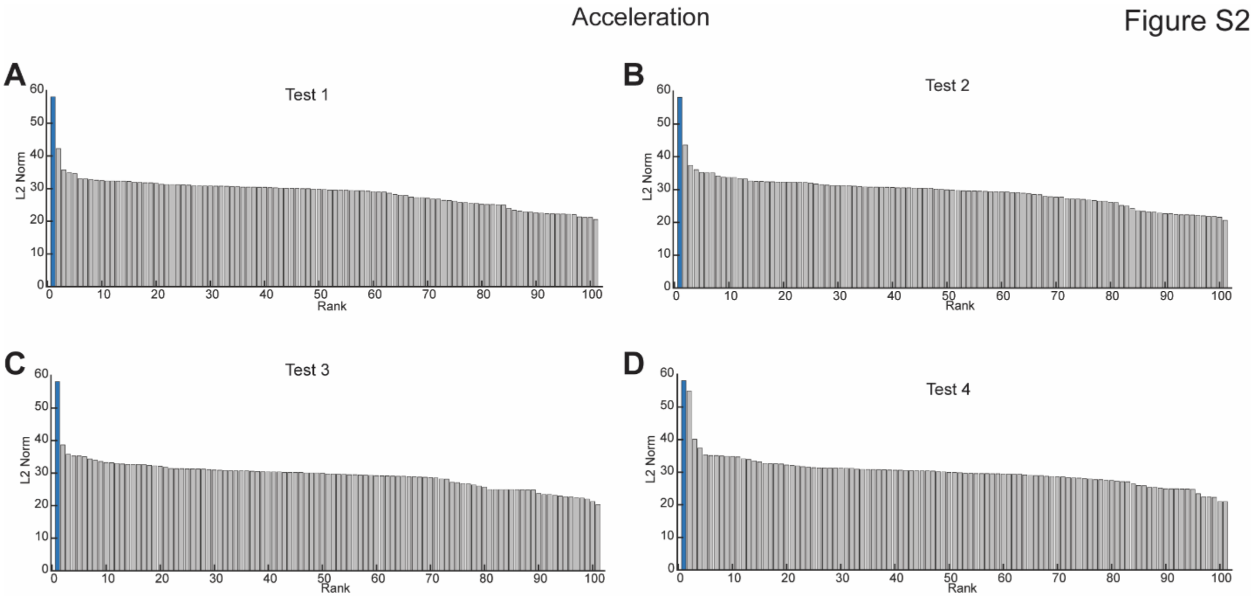


**Figure S2: Comparison of shuffled data significance tests.** Results from comparison of four methods of shuffling data for statistical significance tests of linear filters. (A) Cyclic shuffling of stimulus vector by a random integer. (B) Cyclic shuffling of behavior vector by a random integer. (C) Random permutation of stimulus vector. (D) Random permutation of behavior vector. Bar plots represent the magnitude of filters, computed as the L2 norm, and are plotted in ranked order from highest to lowest magnitude. Colored bar represents appropriately computed filter, gray bars represent filters computers with shuffled data.

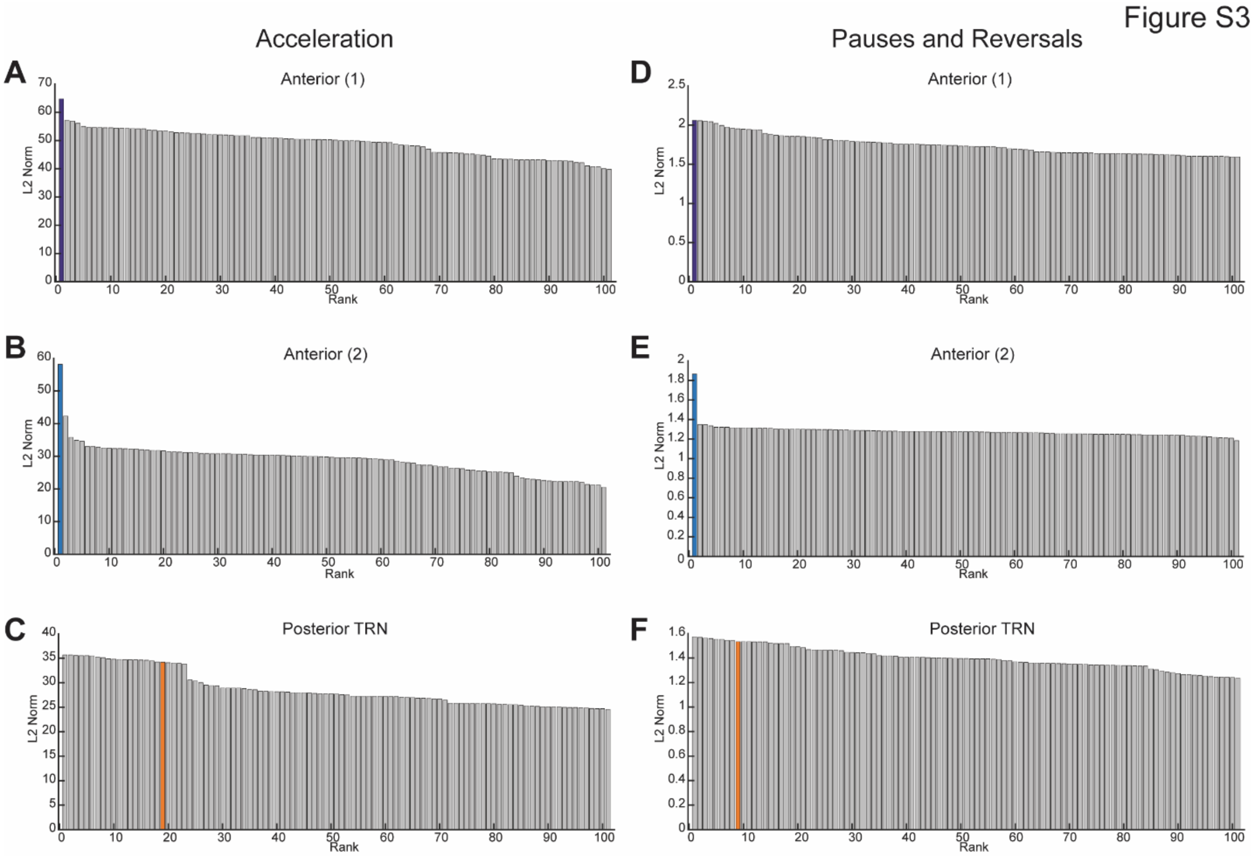


Figure S3: Significance test results for linear filters for TRNs. Results from shuffled data significance tests for linear filters computed for activation of TRNs in Figure 2. (A-C) Significance test results for computed filters for acceleration for anterior TRNs (A,B) and posterior TRNs (C). (D-F) Significance test results for computed filters for pauses and reversals for anterior TRNs (D,E) and posterior TRNs (F). Bar plots represent the magnitude of filters, computed as the L2 norm, and are plotted in ranked order from highest to lowest magnitude. Colored bar represents appropriately computed filter, gray bars represent filters computers with shuffled data.

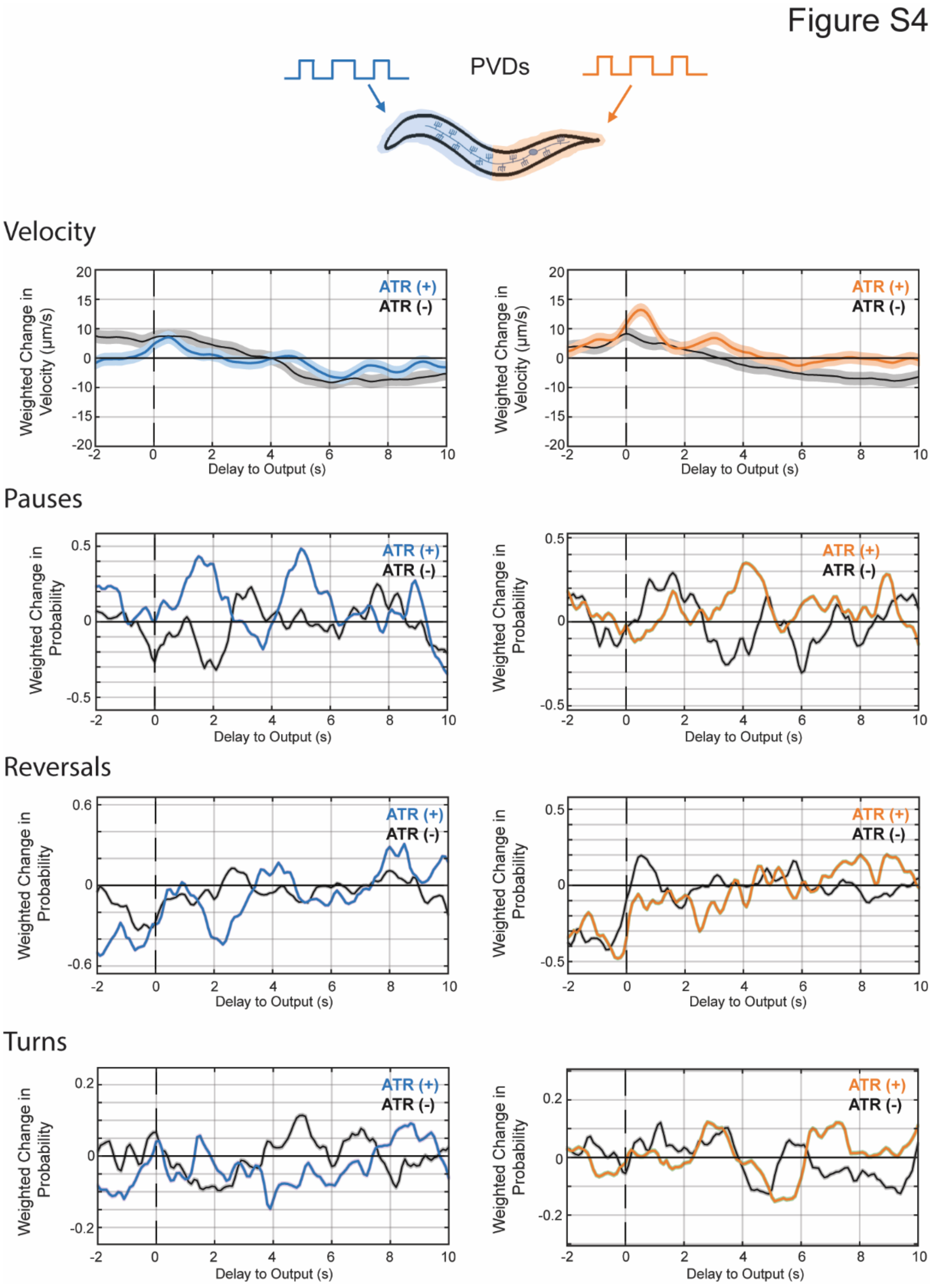


Figure S4: Additional linear filters for PVDs. Linear filters computed for various behaviors when stimulating the anterior (left) and posterior (right) PVDs with an m-sequence signal. Dark line and light shade represent BWA and SEM, respectively. Colored plots represent filters computed from ATR-fed animals, black plots represent filters computed from control (not ATR-fed) animals. Sample sizes are listed Table S1.

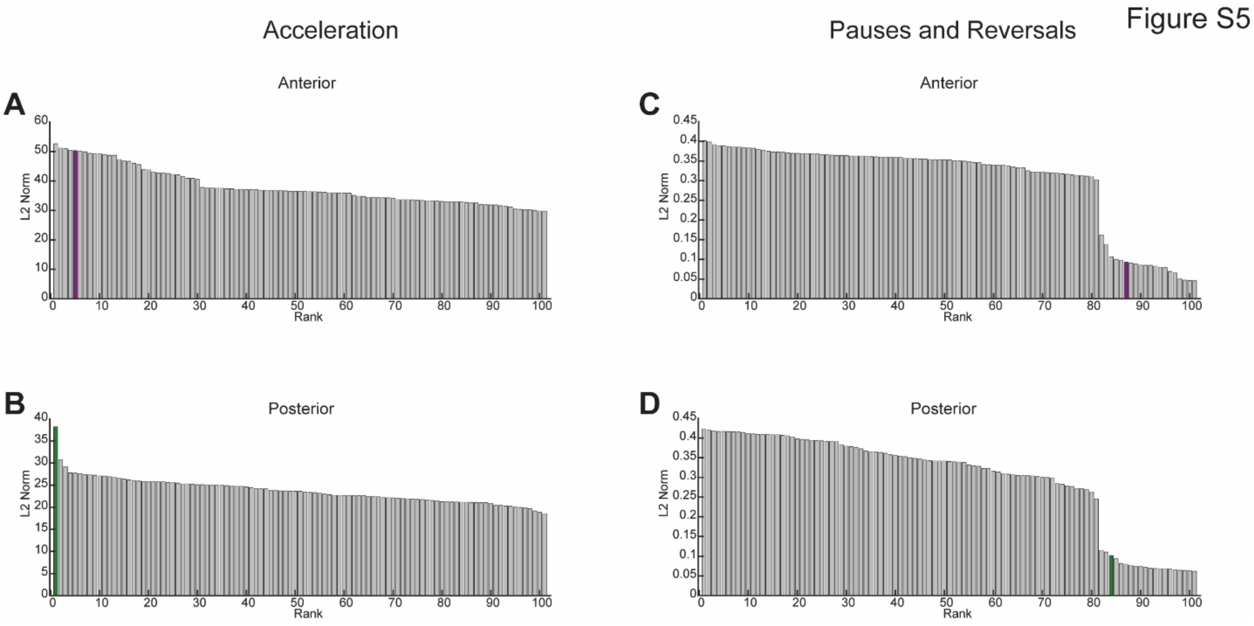


Figure S5: Significance test results for linear filters for PVD. Results from shuffled data significance tests for linear filters computed for activation of PVD in Figure 3. (A,B) Significance test results for computed filters for acceleration for anterior (A) and posterior (B) segments of PVD. (C,D) Significance test results for computed filters for pauses and reversals for anterior (C) and posterior (D) segments of PVD. Bar plots represent the magnitude of filters, computed as the L2 norm, and are plotted in ranked order from highest to lowest magnitude. Colored bar represents appropriately computed filter, gray bars represent filters computers with shuffled data.

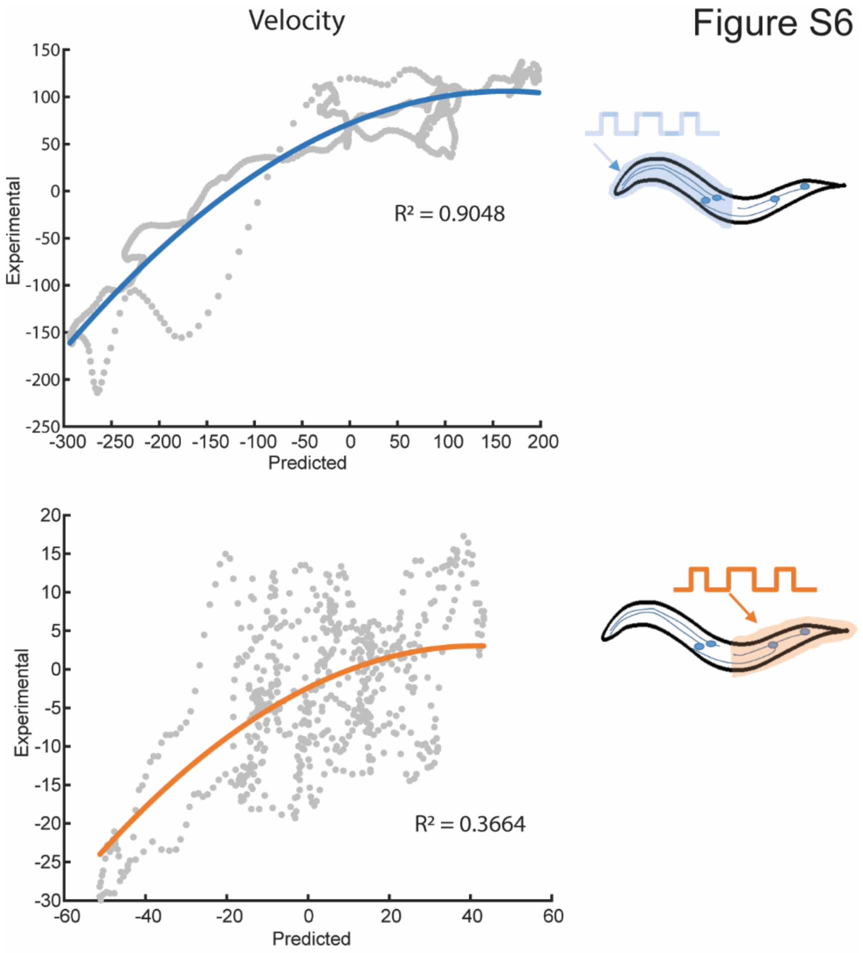


Figure S6: Static nonlinear filters for velocity. Static nonlinear filters fitted for predicted values from the linear filter (x-axis) against experimental values (y-axis) when stimulating the anterior TRNs. Linear filters and experimental values are subsets of data used in Figure 2 (n>1,730 for all conditions). Colored traces represent computed nonlinear filters and gray dots represent independent time-points from measured and predicted values.

Figure S7

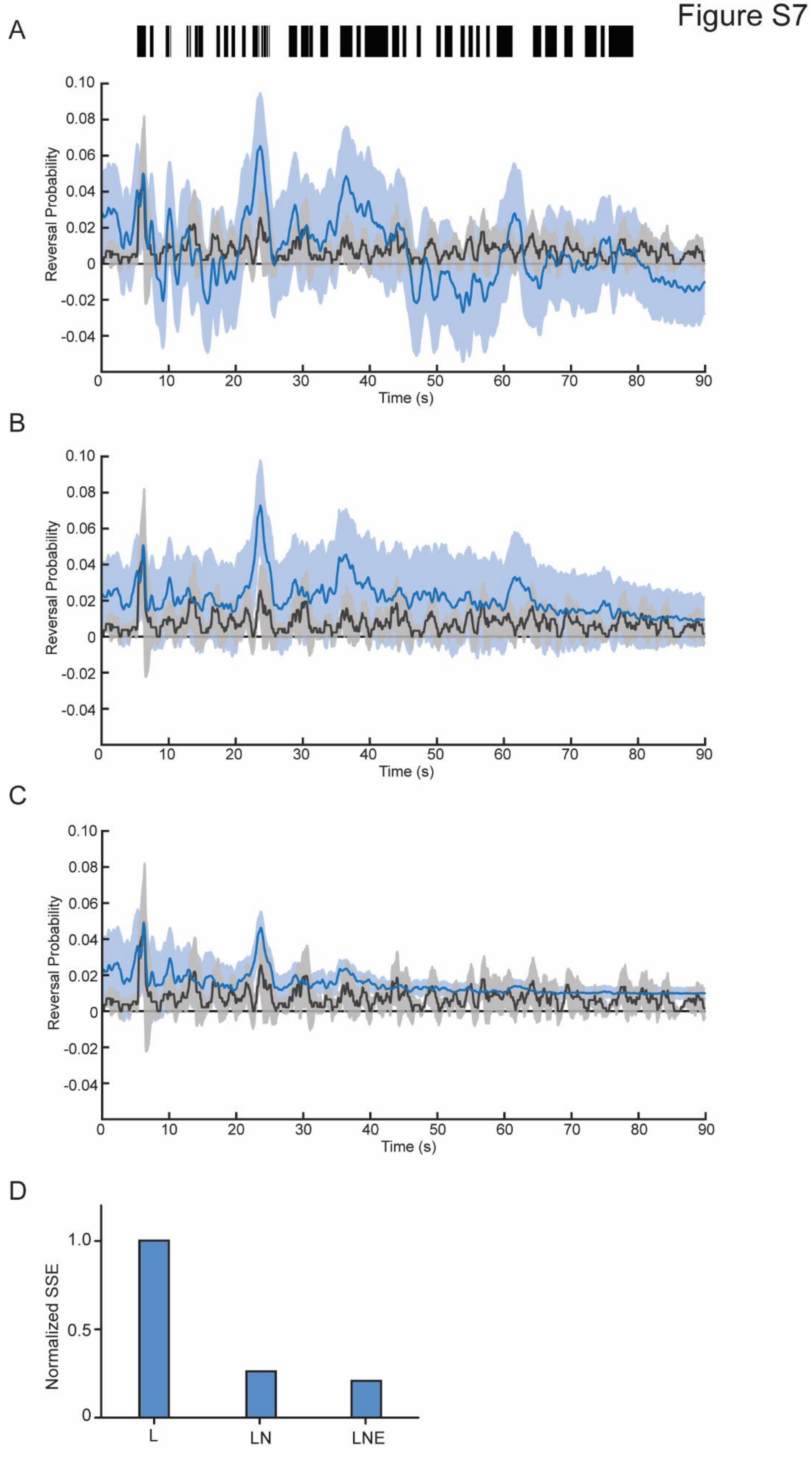

A
Reversal Probability
0.10
0.08
0.06
0.04
0.02
0
-0.02
-0.04
0 10 20 30 40 50 60 70 80 90
Time (s)
B
C
D
Normalized SSE
1.0
0.5
0
L
LN
LNE

Figure S7: Model predictions of reversal initiations. Comparison of model predictions of reversal transitions (blue) and experimental traces (black) when using A) only the linear filter, B) a linear-nonlinear (LN) model, and C) an additional exponential component (LNE). For experimental data, dark line and shade represent average and SEM, respectively (n = 31 animals). For model predictions, dark line represents model prediction and shaded area represents the 95% confidence interval (Methods). Probability of reversal transitions is computed as the average of animals initiating a reversal at that time point. D) Comparison of performance of models, computed as the sum of squared error (SSE) and normalized to the linear model performance value.

Figure S8

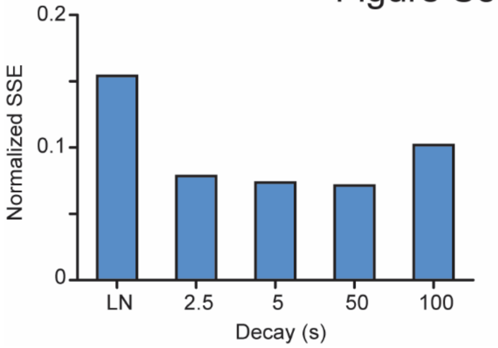


Figure S8: Comparison of decay factors. Comparison of model predictions of velocity for various exponential decay factors. Exponential decays of 2.5s, 5s, 50s, and 100s were tested, with 50s showing the best fit. Performance of models is computed as the sum of squared error (SSE), normalized to the linear model performance value.

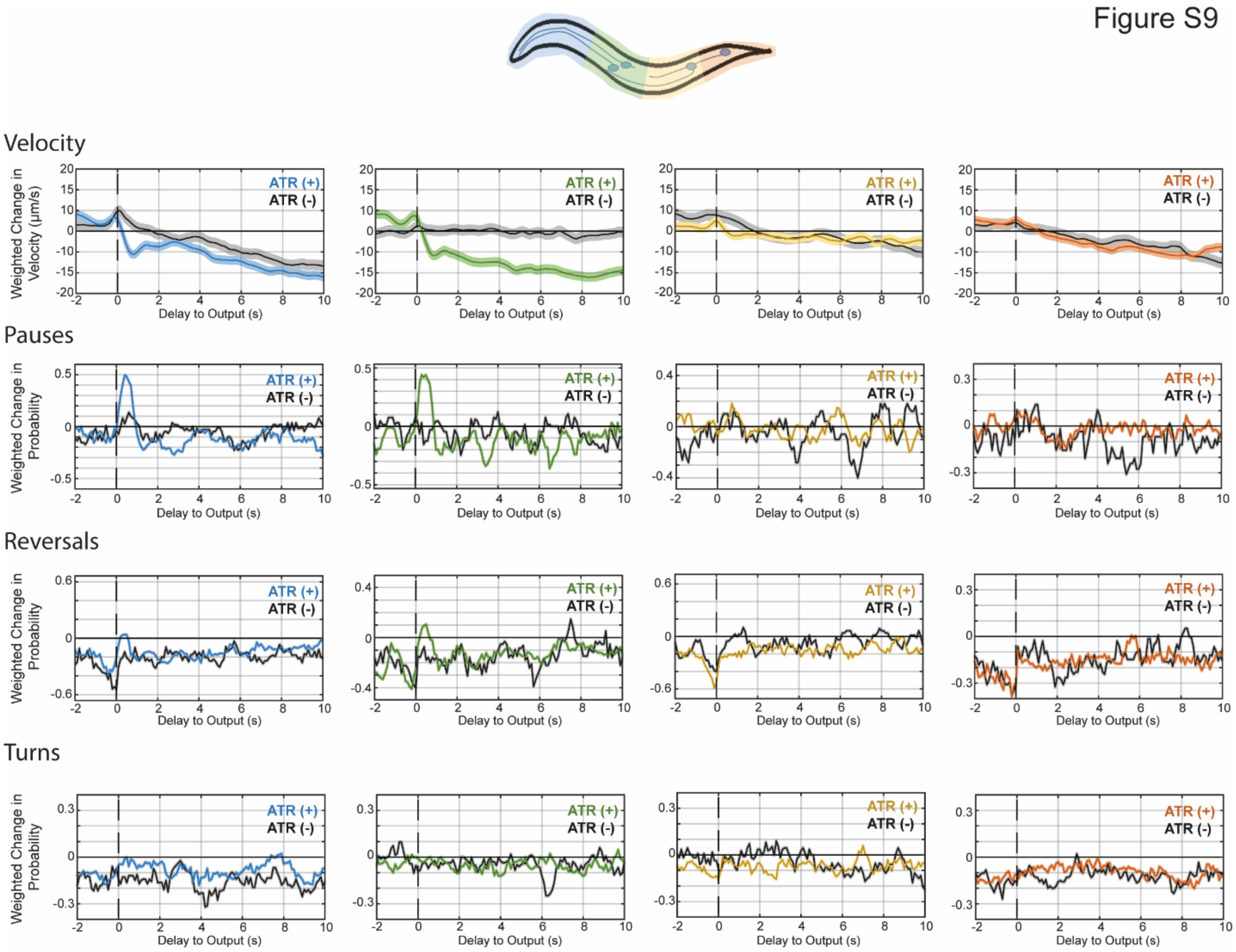


Figure S9: Additional filters for spatially refined analysis of TRNs Linear filters computed for various behaviors when stimulating the most anterior quarter (left), the second-most anterior quarter (second from left), the second-most posterior quarter (second from right), and the most posterior quarter (right) of the TRNs with an m-sequence signal. Dark line and light shade represent BWA and SEM, respectively. Colored plots represent filters computed from ATR-fed animals, black plots represent filters computed from control (not ATR-fed) animals. Sample Sizes are listed Table S1.

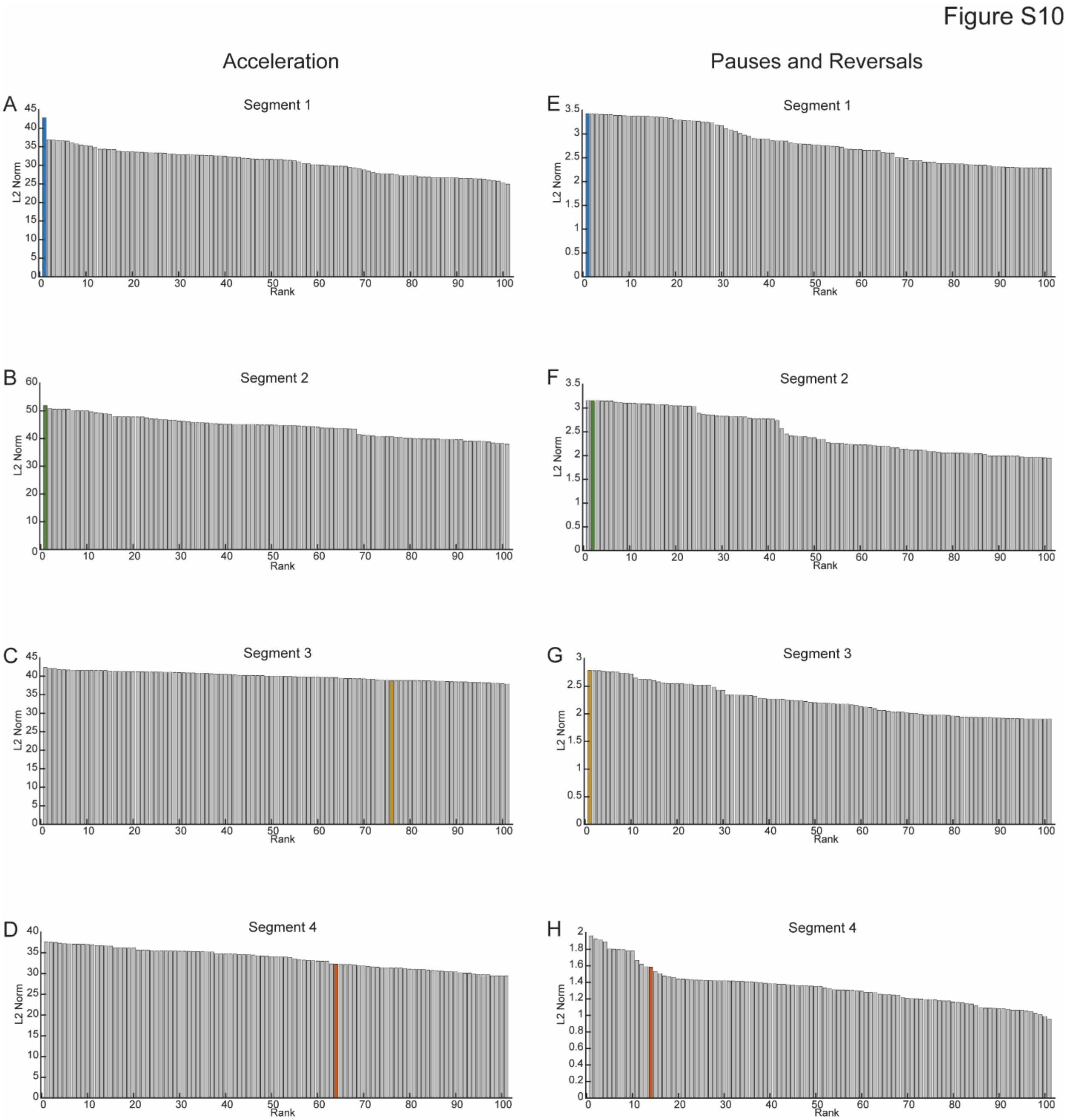


**Figure S10: Significance test results for linear filters for refined TRN analysis.** Results from shuffled data significance tests for linear filters computed for activation of TRNs in Figure 6. (A-D) Significance test results for computed filters for acceleration for most anterior (A), second-most anterior (B), second-most posterior (C), and most posterior (D) segments of TRNs. (E-H) Significance test results for computed filters for pauses and reversals for most anterior (E), second-most anterior (F), second-most posterior (G), and most posterior (H) segments of TRNs. Bar plots represent the magnitude of filters, computed as the L2 norm, and are plotted in ranked order from highest to lowest magnitude. Colored bar represents appropriately computed filter, gray bars represent filters computers with shuffled data.

| Figure | Segment | Behavior | ATR | | nonATR | |
|---|---|---|---|---|---|---|
| | | | Time-Points/Events | Animals | Time-Points/Events | Animals |
| Figure 1D | Anterior | Acceleration | 88031 | 113 | 75618 | 98 |
| Figure 2 | Anterior (1) | Acceleration | 88031 | 113 | 75618 | 98 |
| | | Pauses + Reversals | 1766 | 113 | 1715 | 98 |
| | Anterior (2) | Acceleration | 36597 | 47 | 38238 | 50 |
| | | Pauses + Reversals | 492 | 47 | 422 | 50 |
| | Posterior | Acceleration | 56744 | 61 | 47281 | 49 |
| | | Pauses + Reversals | 667 | 61 | 571 | 49 |
| Figure S1 | Anterior (1) | Velocity | 88025 | 113 | 75618 | 98 |
| | | Pauses | 643 | 113 | 1026 | 98 |
| | | Reversals | 549 | 113 | 163 | 98 |
| | | Turns | 745 | 113 | 876 | 98 |
| | Anterior (2) | Velocity | 36606 | 47 | 38238 | 50 |
| | | Pauses | 156 | 47 | 218 | 50 |
| | | Reversals | 203 | 47 | 37 | 50 |
| | | Turns | 176 | 47 | 242 | 50 |
| | Posterior | Velocity | 56742 | 61 | 47281 | 49 |
| | | Pauses | 197 | 61 | 191 | 49 |
| | | Reversals | 212 | 61 | 133 | 49 |
| | | Turns | 280 | 61 | 272 | 49 |
| Figure 3 | Anterior | Acceleration (max) | 53878 | 61 | 45479 | 51 |
| | | Acceleration (min) | 53816 | 61 | 45404 | 51 |
| | | Pauses + Reversals | 491 | 61 | 555 | 51 |
| | Posterior | Acceleration (max) | 53878 | 61 | 45479 | 51 |
| | | Acceleration (min) | 53807 | 61 | 45464 | 51 |
| | | Pauses + Reversals | 491 | 61 | 555 | 51 |
| Figure S4 | Anterior | Velocity | 53834 | 61 | 45474 | 51 |
| | | Pauses | 98 | 61 | 62 | 51 |
| | | Reversals | 94 | 61 | 161 | 51 |
| | | Turns | 313 | 61 | 340 | 51 |
| | Posterior | Velocity | 53834 | 61 | 45474 | 51 |
| | | Pauses | 98 | 61 | 62 | 51 |
| | | Reversals | 94 | 61 | 161 | 51 |
| | | Turns | 313 | 61 | 340 | 51 |
| Figure 4<br>Figure S5 | Anterior | Acceleration | 600 | 113 | | |
| | | Velocity | 600 | 113 | | |
| | | Pauses + Reversals | 600 | 113 | | |
| | Posterior | Acceleration | 600 | 61 | | |
| | | Velocity | 600 | 61 | | |

| | | | | | | |
|---|---|---|---|---|---|---|
| | | Pauses + Reversals | 600 | 61 | | |
| Figure 6 | Seg1 | Acceleration | 58713 | 67 | 35477 | 40 |
| | | Pauses + Reversals | 538 | 67 | 302 | 40 |
| | Seg2 | Acceleration | 59683 | 69 | 30729 | 35 |
| | | Pauses + Reversals | 434 | 69 | 160 | 35 |
| | Seg3 | Acceleration | 56490 | 66 | 28875 | 33 |
| | | Pauses + Reversals | 366 | 66 | 127 | 33 |
| | Seg4 | Acceleration | 62815 | 71 | 32219 | 36 |
| | | Pauses + Reversals | 545 | 71 | 194 | 36 |
| Figure S9 | Seg1 | Velocity | 58662 | 67 | 35502 | 40 |
| | | Pauses | 129 | 67 | 110 | 40 |
| | | Reversals | 409 | 67 | 192 | 40 |
| | | Turns | 272 | 67 | 134 | 40 |
| | Seg2 | Velocity | 59650 | 69 | 30729 | 35 |
| | | Pauses | 100 | 69 | 65 | 35 |
| | | Reversals | 334 | 69 | 95 | 35 |
| | | Turns | 323 | 69 | 165 | 35 |
| | Seg3 | Velocity | 56480 | 66 | 28876 | 33 |
| | | Pauses | 149 | 66 | 50 | 33 |
| | | Reversals | 217 | 66 | 77 | 33 |
| | | Turns | 251 | 66 | 112 | 33 |
| | Seg4 | Velocity | 62808 | 71 | 32216 | 36 |
| | | Pauses | 224 | 71 | 58 | 36 |
| | | Reversals | 321 | 71 | 136 | 36 |
| | | Turns | 342 | 71 | 145 | 36 |

Table S1: Sample sizes for computed linear filters in figures 1, 2, S1, 3, S4, 4, S5, 6, and S9.
